# Estimating relatedness between malaria parasites

**DOI:** 10.1101/575985

**Authors:** Aimee R. Taylor, Pierre E. Jacob, Daniel E. Neafsey, Caroline O. Buckee

## Abstract

Understanding the relatedness of individuals within or between populations is a common goal in biology. Increasingly, relatedness features in genetic epidemiology studies of pathogens. These studies are relatively new compared to those in humans and other organisms, but are important for designing interventions and understanding pathogen transmission. Only recently have researchers begun to routinely apply relatedness to apicomplexan eukaryotic malaria parasites, and to date have used a range of different approaches on an ad hoc basis. It remains unclear how to compare different studies, therefore, and which measures to use. Here, we systematically compare measures based on identity-by-state and identity-by-descent using a globally diverse data set of malaria parasites, *Plasmodium falciparum* and *Plasmodium vivax*, and provide marker requirements for estimates based on identity-by-descent. We formally show that the informativeness of polyallelic markers for relatedness inference is maximised when alleles are equifrequent. Estimates based on identity-by-state are sensitive to allele frequencies, which vary across populations and by experimental design. For portability across studies, we thus recommend estimates based on identity-by-descent. To generate reliable estimates, we recommend approximately 200 biallelic or 100 polyallelic markers. Confidence intervals illuminate inference across studies based on different sets of markers. These marker requirements, unlike many thus far reported, are immediately applicable to haploid malaria parasites and other haploid eukaryotes. This is the first attempt to provide rigorous analysis of the reliability of, and requirements for, relatedness inference in malaria genetic epidemiology, and will provide a basis for statistically informed prospective study design and surveillance strategies.

## 2 Introduction

Genetic relatedness is a measure of recent shared ancestry (reviewed in [1, 2]). It ranges from zero between two unrelated individuals to one between clones, and in the absence of inbreeding is broken down by recombination [3]. Since the early 20th century, estimates of relatedness have been used across a wide variety of fields: archaeology, agriculture, forensic science, paternity testing, human disease gene mapping, conservation, and ecology [1, 4]. Nevertheless, studies of relatedness are both new and niche in infectious disease molecular epidemiology: new because the field itself is [5], and niche because only a subset of pathogens are eukaryotes, e.g. helmiths and parasitic protoza, which include malaria parasites (reviewed in [6], but without reference to relatedness). Because relatedness is broken down by outbreeding, it can change with each generation [7]. On a population-level, studies of malaria parasite relatedness thus provide a sensitive measure of recent gene flow [8], generating insight on an operationally relevant scale for disease control efforts [9].

Malaria parasites are haploid during the human stage of their life cycle. One measure of relatedness between haploid genotypes is equivalent to the diploid coefficient of inbreeding, defined by Malécot as a probability of identity-by-descent (IBD) [10]. Two alleles are identical by descent (also IBD hereafter) if they are descended from a common ancestor in some ancestral reference population, whose members are assumed unrelated [11, 7, 2].^1^ For pedigrees, the reference is the founder population; more generally, it is a population at some arbitrary time-depth (reviewed in [7, 2]). Two alleles that share the same allelic type are identical by state (IBS) and include those that are both IBD and not IBD [11, 1, 12, 7, 13, 2]. identity-by-state (also IBS hereafter) is observed, whereas IBD is hidden. Though hidden, relatedness based on IBD can be inferred from genetic data. Many estimators exist, some assuming independence between hidden IBD states [1, 11], others not (e.g. [14] and subsequent models - see [15] and references therein). Those assuming independence have fewer parameters but impaired power in the presence of dependence [16]. Estimators that do not assume independence are often based on hidden Markov models (HMMs, reviewed in [17]). Measures of relatedness used in studies of malaria include those estimated under HMMs (hmmIBD [18], used in e.g. [19, 20, 21, 22]; isoRelate [23], extension of XIBD [24]; DEploidIBD [25], extension of DEploid [26]). Measures based on IBS (e.g. proportions of alleles shared or counts of allele differences) require only simple calculation and are thus popular also as a proxy for measuring relatedness between malaria parasites (e.g. [27, 28, 19, 29, 30, 31, 32]).

Despite many malaria genetic epidemiology studies using IBD and IBS based analyses, there are few systematic comparisons applicable to malaria studies. Questions also remain about optimal marker requirements for pairwise relatedness inference. To enable comparison between studies of malaria epidemiology, we compare and assess measures based on IBD and IBS using simulated data; various data sets of *Plasmodium falciparum*, the parasite responsible for the most deadly type of human malaria; and a data set of *Plasmodium vivax*, the parasite most commonly responsible for recurrent malaria. We use a model framework encompassing two simple models assuming independence and not. It is an error-modified version of [14] and thus at the core of many probabilistic IBD models (see review in [15]), including those specifically designed for comparison across malaria parasites [18, 23]. To guide future relatedness studies of monoclonal haploid malaria parasite samples and haploid eukaryotes more generally, we explore marker count and number of alleles for relatedness inference with specified error. Simulated data illustrate how IBS is sensitive to marker panels, making conclusions non-portable across studies. Concrete recommendations on marker requirements depend on specified error. Increasing the number of alleles per marker genotyped reduces error, especially when markers are few, but with diminishing returns.

## 3 Methods and Theory

### 3.1 Relatedness

For the purpose of this study, relatedness *r* is defined as the probability that, at any locus on the genome, the allele sampled from one individual is IBD to the allele sampled from the other individual. This is referred to as the pointwise pairwise probability of IBD in [15]. We denote by *m* the number of genotyped markers. Each of them has a locus on the genome. We index these loci by *t* = 1*, …, m*. We denote by *c*_*t*_ the index of the chromosome of the *t*-th locus, and by *p*_*t*_ its position on that chromosome. For two indices *t*_1_*, t*_2_ with *t*_1_ < *t*_2_, we either have 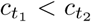, or 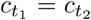 and 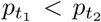. For the *t*-th locus we denote by IBD_*t*_ the binary variable indicating whether the two individuals are IBD at that locus; IBD_*t*_ = 1 indicates IBD, otherwise IBD_*t*_ = 0.^2^ We assume that *r*, the marginal probability that IBD_*t*_ = 1, is constant across the genome:

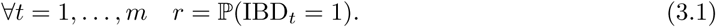

The sequence (IBD_*t*_)_*t*=1*,…,m*_ could be made of independent Bernoulli variables with parameter *r*, or more generally a Markov chain with a Bernoulli invariant distribution with parameter *r*. For the Markov chain model, we write the transition probabilities at locus *t*,

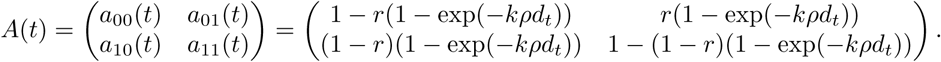

In the above, *a*_*jℓ;*_(*t*) refers to the probability of IBD_*t*_ = ℓ; given that IBD_*t*-1_ = *j*, *d*_*t*_ denotes a genetic distance in base pairs (bp) between sites *t -* 1 and *t* (i.e. all markers are treated as point polymorphisms). If the locus *t -* 1 and *t* are on different chromosomes (*c*_*t-*1_≠*c*_*t*_) the distance is set to +*∞*; in that case the variables IBD_*t*-1_ and IBD_*t*_ are independent. The value *k >* 0 parameterizes the switching rate of the Markov chain and *ρ* is a constant equal to the recombination rate, assumed known and fixed across both haploid genotypes with value 7.4*×*10^*−7*^M bp^*−1*^ for *P. falciparum* parasites [33].

We now describe how the model connects the quantity of interest *r* to the data. At each locus *t*, we assume that the set of possible alleles is denoted by 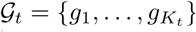, where *K*_*t*_ ≥ 2 denotes the cardinality of 𝒢_*t*_ (allelic richness of the *t*-th marker). For individuals *i, j* in the population and at locus *t* we observe the pair 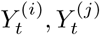 ∈ 𝒢_*t*_. We assume that alleles occur with frequencies 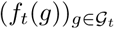, with *f*_*t*_(*g*) ≥ 0 for all *g ∈𝒢*_*t*_ and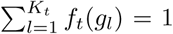. The data comprise 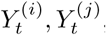, the distances (*d*_*t*_) and the frequencies 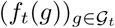 at *m* loci. A simple observation model relates the data to IBD_*t*_ by assuming that, if IBD_*t*_ = 0, then 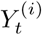and 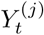 are independent categorical variables taking values in 𝒢_*t*_ with probabilities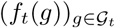. If IBD_*t*_ = 1, then 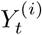 is such a categorical variable and 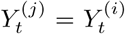 with probability one. A more realistic model accounting for observation error is described in Appendix B.

Combining the Markov model for (IBD_*t*_) with an observation model as above leads to a hidden Markov model (Figure 1) with a likelihood function (*r, k*) ↦ 𝓛_1:*m*_(*r, k*). Note that, as mentioned in the introduction, this model is essentially an error-modified version of [14], and thus at the core of the many subsequent probabilistic IBD models (see [15]), including all those specifically designed for comparison across haploid malaria parasites [18, 23]. An independence model can be retrieved by e.g. setting all distances to +*∞*. In either case we can maximize the likelihood over the parameter space and we denote by 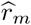 the maximum likelihood estimator of *r*, that can be computed using numerical optimization.

**Figure 1:**
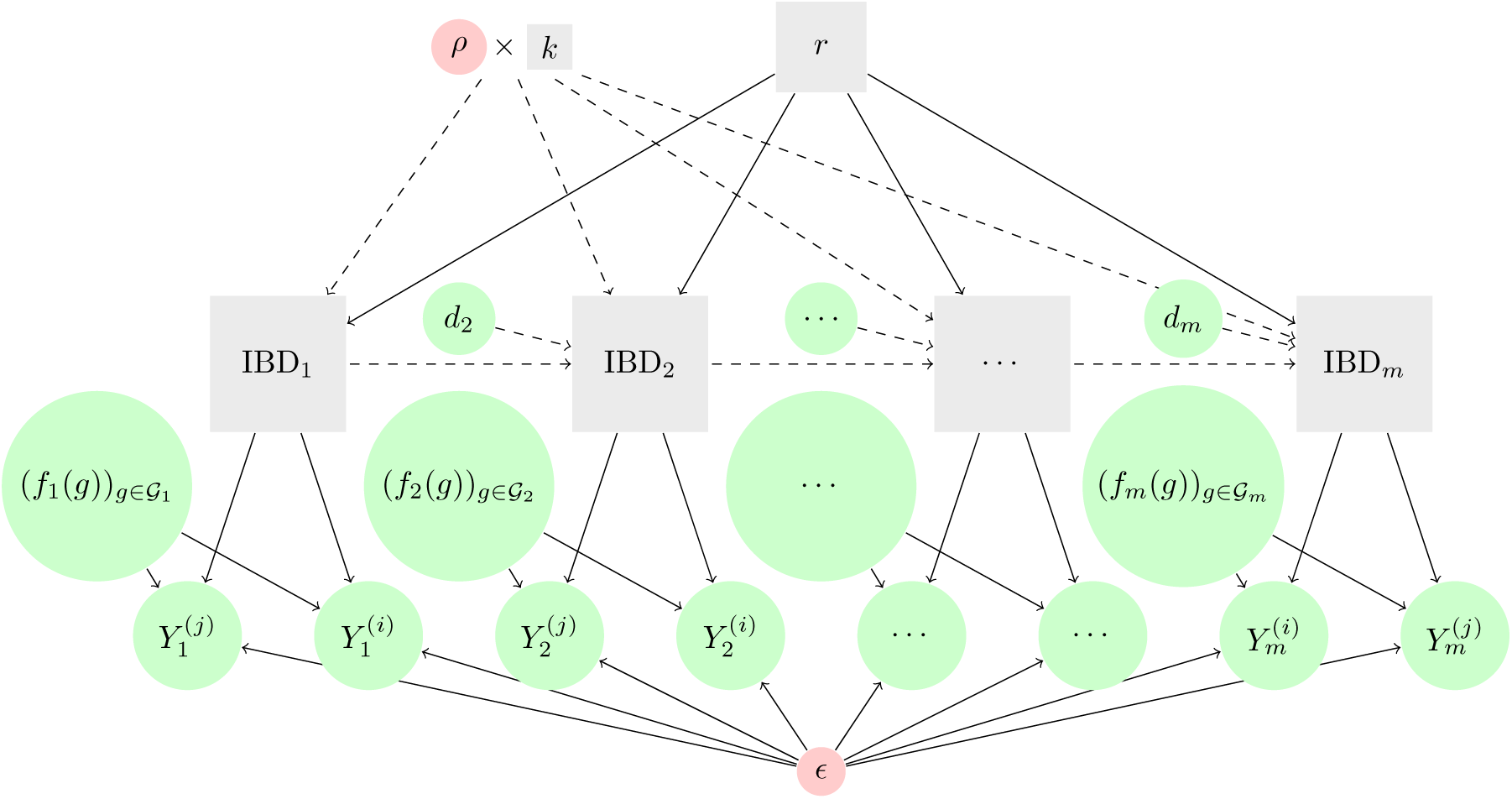
Models relating genetic data to genetic relatedness. Input data are depicted by green circles. They include genotype calls, 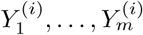 and^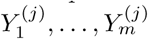^, allele frequencies, 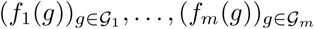, and distances between genotyped markers, *d*_2_*, …, d*_*m*_. Parameters considered fixed (the genotyping error, ϵ, and the constant, *ρ*) are depicted by red circles. Unobserved quantities are depicted by gray squares. They include the hidden IBD states, IBD_1_, …, IBD_*m*_, and the estimands (genetic relatedness, *r*, and *k*). Solid arrows depict dependencies under both the independence model and the HMM. Dashed arrows depict dependencies under the HMM only. Distances, *ρ* and *k* feature in the HMM only.

Under some assumptions on the data-generating process, the maximum likelihood estimate 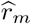 could be shown to be consistent for *r* as the sample size *m* goes to infinity. However these asymptotic considerations are intricate in the present setting of Mendelian sampling [34]. Indeed, the degree of dependencies between successive observations increases with the sample size *m*: the more sites are sampled, the closer to one another they become. This departs from the standard asymptotic setting, where the observations are not increasingly dependent as *m → ∞* [35, 36, 37]; this is discussed in more details in Appendix B.

Without standard asymptotic results, such as the asymptotic normality of the maximum likelihood estimator, we do not have a simple formula relating the sample size *m* to the variance of the estimator _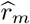_, which would have been useful for sample size determination. The distribution of the 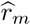 can still be approximately normal because the log-likelihood can be approximately quadratic [38]. If that is the case, confidence intervals can be obtained through the second derivative of the log-likelihood at the MLE. The present setting poses an additional difficulty since the MLE can be located on the boundary of the parameter space, e.g. 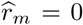 or 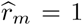 This suggests that the distribution of the MLE might not always be normal [39]. We therefore rely on the parametric bootstrap [Chapter 9 of 40] to construct confidence intervals around 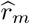 (note that we cannot use the nonparametric bootstrap since we cannot sample positions with replacement). Unless otherwise stated, we use 500 bootstrap draws throughout. Note that even when assuming the absence of genotyping error, if 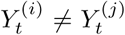 for some *t* = 1*, …, m*, the confidence interval around 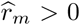 can contain one, since data simulated with 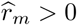 may have identical genotype calls for all *t* = 1*, …, m* (especially if *m* is small), which leads to a bootstrapped estimate of *r* equal to one.

Under the model framework (Figure 1) no explicit mention is made of ancestors, be they most recent or at some arbitrary time-depth. In the introduction we refer to IBD relative to either some reference population or some small segment length (‘recombination-sense’ [2]). Akin to many existing IBD models (see catalogue in [15, 41, 42]) and some related imputation methods (see catelogue in [43, 44, 45]), IBD segments can be estimated within the HMM framework (e.g. using the most likely path of hidden states [17], posterior probabilities of IBD at each marker position [42, 46], or a posterior predictive draw of IBD segments [46]). These segments underpin many applications from disease mapping (e.g. [47]) to *P. falciparum* selection detection [23]. They can also be used to generate a recombination-sense IBD estimate [2]. However, like [15] we do not tune the parameter which relates to segment length, *k*. As such the pointwise estimate _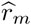_, averages over all IBD segments, however small, and thus is liable to reflect some antecedent population-level relatedness, i.e. linkage disequilibrium (LD) [48]).

### 3.2 Fraction IBS

For a pair of samples *i* and *j*, we define the fraction IBS as a proportion of *m* markers that are identical across both samples,

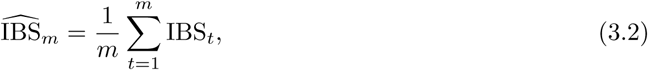

where IBS_*t*_ = 1 if 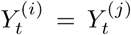 and zero otherwise. The fraction IBS can also be derived from counts of marker differences; e.g. 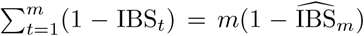 where 1 − IBS_*t*_ = 1 denotes a marker difference (equation (3.2) rearranged). Note that 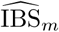 is the haploid equivalent of the ‘allele-sharing coefficient’ referred to in [2].

We can relate the IBS estimator (equation (3.2)) to *r* by specifying a relationship between IBD_*t*_ and IBS_*t*_. For illustration, in the case with no genotyping error, the expectation of IBS_*t*_ is the following linear function of relatedness (derivation, equation (A.1)),

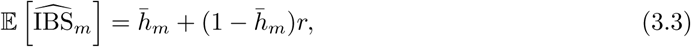

Where

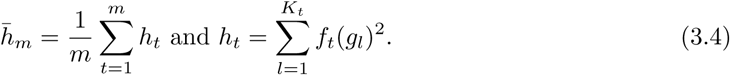

Here *h*_*t*_ and 1*-h*_*t*_ are equivalent to Nei’s ‘gene identity’ and ‘gene diversity’, respectively [49]. Considering an outbred diploid, these terms equate to homozygosity and heterozygosity, respectively [49].

Equation (3.3) might suggest that 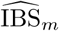 could converge to 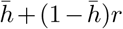 (where 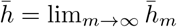) as the number of markers *m* goes to infinity, under assumptions on the data-generating process such as independent loci; see section A.2. Under this setup, the estimator 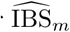 would not be consistent for *r*, but could be corrected; see Appendix A. In the Results section we numerically demonstrate how equation (3.3) is a problematic estimator of *r* using simulated and real data (see details below).

### 3.3 Plasmodium data

Throughout, we illustrate results using *Plasmodium* and simulated data. *P. falciparum* data comprise biallelic (i.e. *K*_*t*_ = 2 ∀*t* = 1*, …, m*) single nucleotide polymorphism (SNP) data from monoclonal *P. falciparum* samples (Table 1). All data are published [50, 51, 29, 30, 52, 8]. They were obtained either from sparse genome-wide panels of select markers, called barcodes, or from dense whole genome sequencing (WGS) data sets (reviewed in [53]); full details of sample collection and data generation can be found via the citations above and references therein. Additional steps we took to process the data are as follows.

**Table 1:** A summary of globally diverse data sets of monoclonal *P. falciparum* samples. All data are published. Full details of sample collection and data generation can be found via the citations above and references therein. Additional steps we took to process the data for use in this study are described in section 3.3. For each processed data set, *n* denotes the number of monoclonal *P. falciparum* samples; *m*_max_ denotes the maximum number of successfully genotyped SNPs per sample; 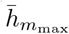 denotes the expected homozygosity (equation (3.4)) averaged over *m*_max_ SNPs; and 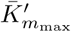 denotes the average effective cardinality (defined below, equation (3.8)).

| Data set and citation/s | Collection region and years | $n$ | $m_{\max}$ | $\bar{h}_{m_{\max}}$ | $\bar{K}'_{m_{\max}}$ |
| --- | --- | --- | --- | --- | --- |
| Colombia [50] | Colombian Pacific region, 1993-2007 | 325 | 250 | 0.66 | 1.57 |
| Thailand 93-SNP [51, 8] | Thailand-Myanmar border, 2001-2010 | 1173 | 93 | 0.57 | 1.77 |
| Thailand WGS [52, 8] | Thailand-Myanmar border, 2001-2014 | 178 | 34911 | 0.89 | 1.17 |
| The Gambia [29] | Kombo coastal districts, 2007–2008 | 71 | 31 | 0.77 | 1.37 |
| Kilifi [29] | Coastal Kenya, 1998-2010 | 628 | 127 | 0.87 | 1.19 |
| Western Kenya [30] | Western Kenya, 2008-2010 | 182 | 59 | 0.73 | 1.43 |

Besides mapping SNP positions to the *P. falciparum* 3d7 v3 reference genome and recoding heteroallelic calls as missing (since all samples with fewer than 10 heteroallelic SNP calls were classified monoclonal by [50]), we did not post-process the Colombian data in any way. Thailand 93-SNP and WGS samples were used exactly as described in [8]. Data derived from [29, 30] (i.e. all African data) were processed using steps described in ‘Sample and SNP cut-off selection criteria’ of [29]. In addition, we removed samples with duplicate SNP calls; removed samples classified as not monoclonal using a *≤*5% heteroallelic SNP call rate to classify monoclonal samples, akin to [51]; and, among monoclonal samples, treated heteroallelic SNP calls as missing and removed monomorphic SNPs.

For each processed data set of monoclonal *P. falciparum* samples, allele frequencies were estimated by simple proportions: 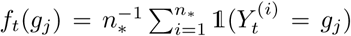 for all *j* = 1, 2 and each locus *t*, where *n*_*_ *≤ n* denotes the number of monoclonal parasite samples whose data were not missing at the *t*-th locus. Minor allele frequencies (i.e. min(*f*_*t*_(*g*_1_)*, f*_*t*_(*g*_2_))*∀t* = 1*, … m*) vary considerably by design (i.e. different marker panels) and due to variation among parasite populations over space and time (Figure 2).

**Figure 2:**
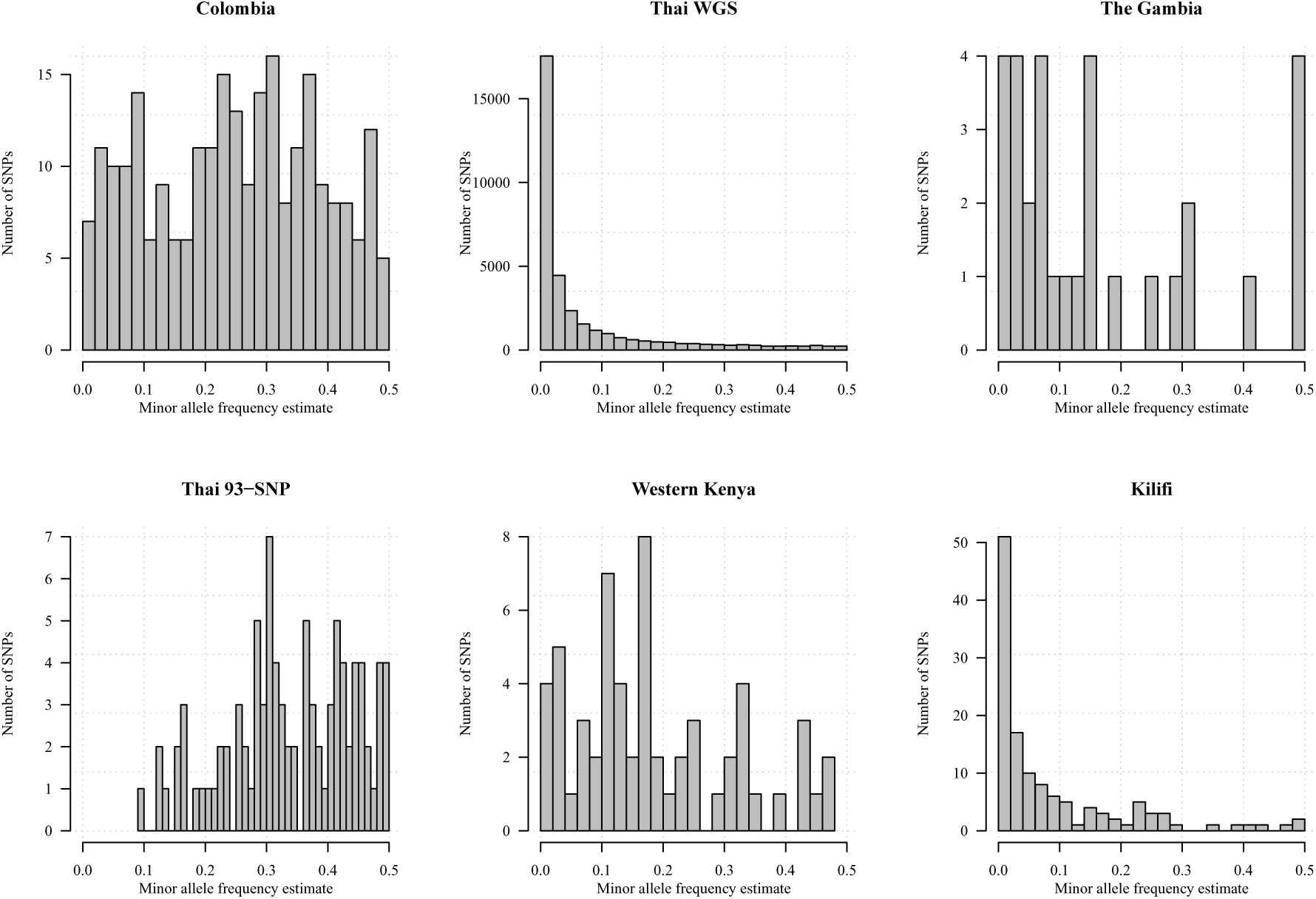
Minor allele frequency estimates from monoclonal *P. falciparum* data sets (Table 1).

In addition to the aforementioned *P. falciparum* data, we generated results for a single *P. vivax* data set, freely available online [54]. The *P. vivax* data were collected between 2010 and 2014 from two clinical trails on the Thailand-Myanmar border [55, 56]. The data were genotyped at three to nine highly polyallelic microsatellites (MS). In this study, we analyse samples genotyped at nine MSs that have no evidence of multiclonality (detection of two or more alleles at one or more MS). We estimate relatedness between pairs of samples from *n* = 204 different people, selecting one episode per person uniformly at random from all episodes per person. We use the allele frequencies reported in [54]. They have average expected homozygosity 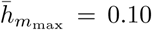 (equation (3.4)) and average effective cardinality (defined below, equation (3.8)) 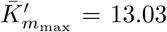..Since there are only nine markers, we analyse these data under the independence model.

### 3.4 Simulated data

#### 3.4.1 Biallelic markers

Unless otherwise stated, biallelic marker data (i.e. data with *K*_*t*_ = 2 ∀*t* = 1*, …, m*) were simulated under the HMM with *ε* = 0.001 using marker loci positions and allele frequency estimates sampled from the Thailand WGS data set. Positions were sampled uniformly at random. Frequencies were sampled separately using one of two approaches: either they were sampled uniformly at random, or, to compensate for the skew towards rare alleles in WGS data set, frequencies were sampled separately with probability proportional to minor allele frequency estimates.

#### 3.4.2 Polyallelic markers

Polyallelic marker data (i.e. data with *K*_*t*_ > 2 *∀t* = 1*, …, m*) were simulated under the HMM with *ε* = 0.001 using marker loci positions sampled uniformly at random from the Thailand WGS data set and allele frequency estimates sampled from a Dirichlet distribution. We used a Dirichlet parameter vector equal to 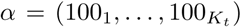 to generate frequencies such that alleles are approximately equifrequent, and a concentration parameter vector equal to 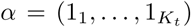 to generate frequencies that are uniform over the *K*_*t*_ − 1 simplex. The former approach generates ideal frequencies (see below), while the latter generates frequencies that for *K*_*t*_ > 2 are increasingly skewed towards rare alleles, thus more representative of real frequency spectra.

### 3.5 Marker requirements for prospective relatedness inference

#### 3.5.1 Biallelic markers

For a set of parameters (i.e. number of markers *m*, relatedness *r*, switch rate parameter *k*) we simulate 1000 pairs of haploid genotype calls and for each pair compute 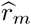. We compute the root mean squared error (RMSE) by taking the square root of the average of the squared difference between 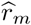 and *r*. From the RMSEs computed for different sample sizes *m*, we derive the number of markers required for the RMSE to be below a specified value. Unless otherwise stated we use *k* = 12 where fixed, the mean estimate of *k* for 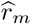 ∈ (0.475,0.525) from the Thailand WGS data set.

Comparison between 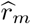 and *r* differs from that between 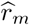 and 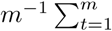 IBD, which is referred to as the ‘realised relatedness’ in [2]. The former has the advantage of revealing RMSE due to the finite length of the genome (i.e. Mendelian sampling [34]), while at the same time revealing the excess, and thus theoretically avoidable, error due to marker limitations.

#### 3.5.2 Polyallelic markers

To explore marker requirements for relatedness inference using polyallelic markers we first consider the impact of increasing *K*_*t*_ beyond two at a single locus. For a given *K*_*t*_, we measure the informativeness of a set of allele frequencies via the corresponding Fisher information matrix; this can in turn be related to the precision of the maximum likelihood estimator if the log-likelihood is approximately quadratic. We define 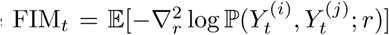, where the expectation is with respect to 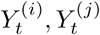 given *r* and the allele frequencies, and where we consider an observation model without genotyping error for simplicity; the sign 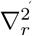 stands for the second order derivative with respect to *r*. The Fisher information matrix FIM_*t*_ depends on the allele frequencies 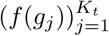 and on *r*:

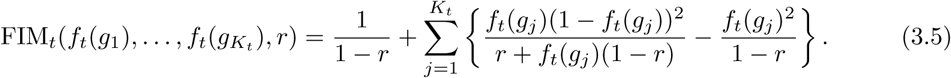

We show that, for any *K*_*t*_ and *r*, it is maximized over all 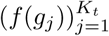 by 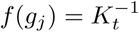 for all *j*, i.e. by equifrequent alleles. This is in agreement with the aforementioned long-established result that markers with high minor allele frequencies are preferable for relatedness inference [57]. A proof is provided in Appendix B.3.4. When alleles are equifrequent we obtain

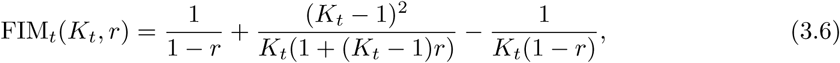

which is an increasing function of *K*_*t*_ such that 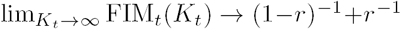. Equation (3.6) describes the precision of the MLE, assuming that the log-likelihood is approximately quadratic, that *K*_*t*_ is the same at each locus and the allele frequencies are equifrequent.

To explore the relative gain of increasing *K*_*t*_ > 2 we calculate the multiplicative increase in FIM_*t*_(*K*_*t*_ ≥ 2*, r*) relative to FIM_*t*_(*K*_*t*_ = 2*, r*) (Figure 3, left). The informativeness of *K*_*t*_ = 15 is between approximately two and seven times that of *K*_*t*_ = 2, with increasing returns as *r* approaches zero. However the justification of the FIM as a measure of precision breaks at the boundary of the parameter space. Regardless of *r*, the biggest increase is obtained upon increasing *K*_*t*_ from 2 to 3 with diminishing returns thereafter. The plot on the right of Figure 3 shows multiplicative increase in precision as a function of effective cardinality,

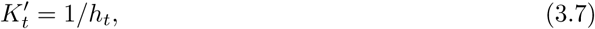

which can be interpreted as the non-integer number of equifrequent alleles that would give rise to the same *h*_*t*_ as that based on the allele frequencies 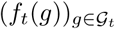 (equation (3.4)). For example, 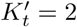 is the effective cardinality of an ideal biallelic SNP, whereas 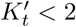 is the effective cardinality of a realistic biallelic SNP. Precision increases with 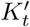 as it does with *K*_*t*_.

**Figure 3:**
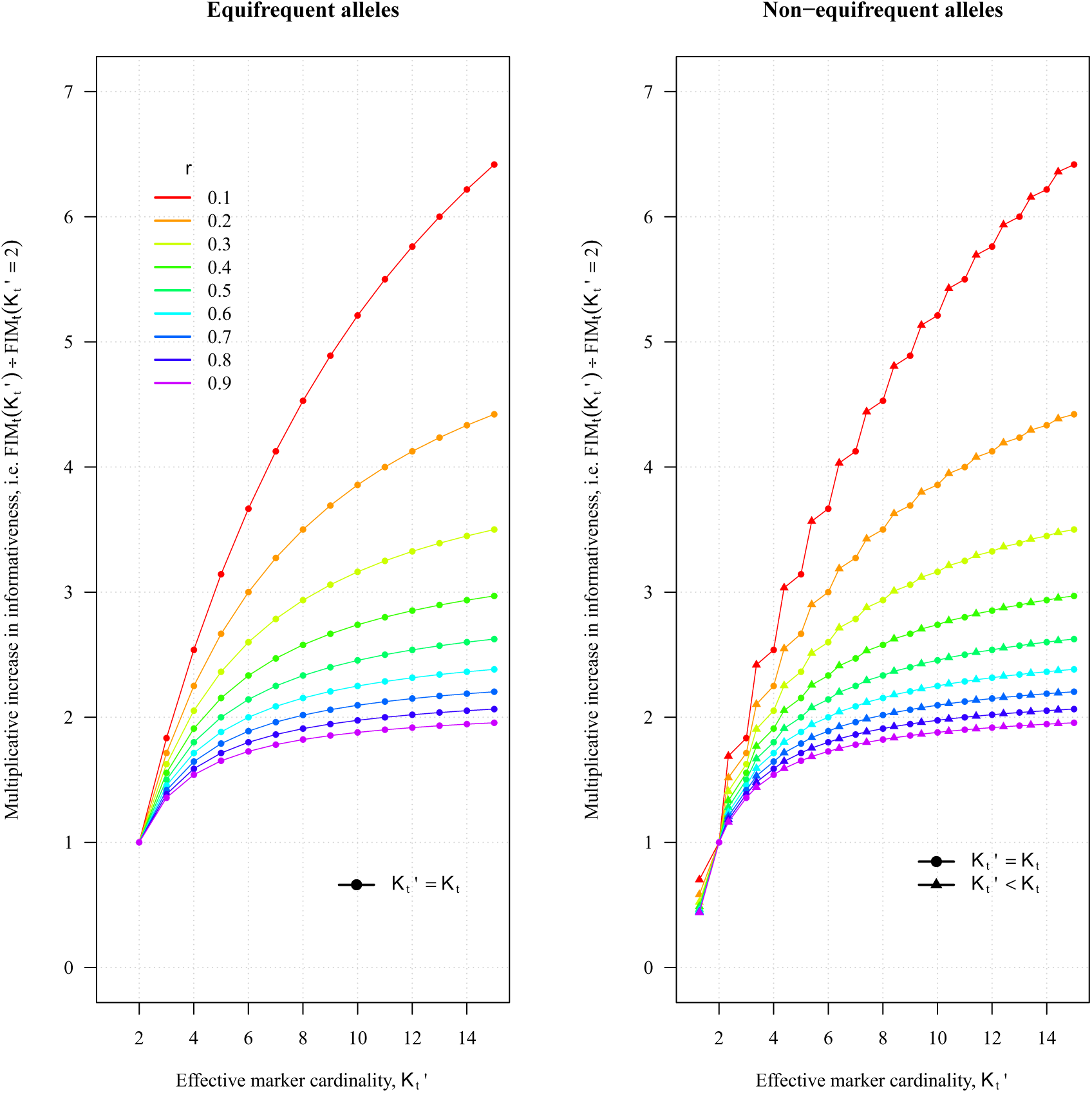
Multiplicative increase in the precision of the MLE with marker cardinality. The left plot shows the multiplicative increase for equifrequent alleles according to equation (3.6). The right plot shows the multiplicative increase with 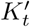, where precision was calculated according to equation (3.5) with either *f*_*t*_(*g*_*i*_) = 1*/K*_*t*_ *∀ i* = 1, …, *K*_*t*_ (dots) or *f*_*t*_(*g*_1_) = 1.75*/K*_*t*_ and *f*_*t*_(*g*_*i*_) = (1−*f*_*t*_*(g*_1_))/ *Kt -* 1) *∀ i* = 2, …, *K*_*t*_ such that 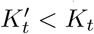 (triangles).

To explore the trade-off between increasing *m* and increasing *K*_*t*_ for a set of parameters (i.e. various *m*, *K*_*t*_, α and *r*, and *k* = 12), we simulate 1000 pairs of haploid genotype calls, generate 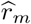 for each pair and calculate the RMSE. For simplicity, for a given *m*, we assume all markers have the same *K*_*t*_. To compare on a common scale numerical results for makers with and without equifrequent alleles, we use the average effective cardinality:

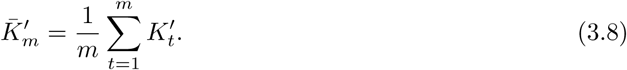

Since 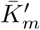 is approximately the same for all *m*, to explore the trade-off between increasing *m* and increasing *K*_*t*_, we average the effective cardinality over all *m* ∈ {24, 96, 192, 288, 384, 480},

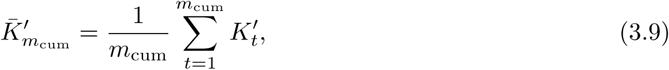

where *m*_cum_ = 24 + 96 + *…* + 480.

### 3.6 Data and code availability

All data used in this study are either simulated or published previously. Additional steps we took to process the data are described in section 3.3. The processed data and code necessary for confirming the conclusions of the article are available at github.com/artaylor85/PlasmodiumRelatedness.

## 4. Results

This section concerns the estimation of *r* as defined above and is arranged as follows. First we consider the genomic fraction IBS and show how it is problematic as an estimator of *r*. Second, we discuss *r* estimated using *Plasmodium* data, and provide marker requirements based on simulated data with biallelic and polyallelic markers.

### 4.1 Fraction IBS

Although 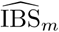 might not satisfy favorable statistical properties as an estimator of *r*, its expectation is indeed related to *r* (equation (3.3)). As such, many studies have recovered meaningful trends in *r* with respect to epidemiological covariates (e.g. geographic distance) using measures related to 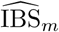 [29, 30, 32]. However, since 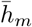 is a function of the allele frequencies (equation (3.4)), so too is 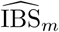. This is equivalent to the dependence on MAFs of the allele-sharing coefficients reviewed in [2]. It means that quantitative trends in 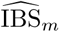 (e.g. regression coefficients) and absolute values of 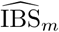 are only comparable across data on markers whose allele frequencies are the same [32]. This is a limitation given that frequencies almost always differ across data sets (e.g. Figure 2).

To illustrate the effect of differing allele frequencies, we generated 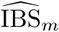 for data simulated using allele frequency estimates from published data sets (Figure 4, top). The plot illustrates two notable results. First, for all biallelic marker data sets, the 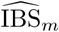 distribution is far from 0.5, the value of *r* used to simulate the data. The expected difference between 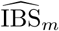 and *r* is a linearly decreasing function of 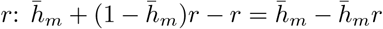. As such, we would expect to see bigger and smaller distances between 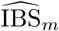 distributions and the data generating *r* were data simulated using *r <* 0.5 and *r >* 0.5, respectively; when *r* = 1 there is no difference. Second, the locations of the 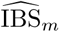 distributions vary considerably across data sets, centering around 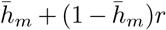, which varies due to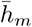, since *r* = 0.5 throughout. This variability, despite all parasite pairs having been simulated with *r* = 0.5, renders absolute values of 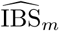 non-portable across data sets. In contrast, distributions of estimates of relatedness based on 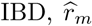, all centre around *r* = 0.5 (Figure 4, bottom). To single out the effect of frequencies, we fixed all parameters besides frequencies across the data sets, including the number of markers (*m* = 59) and their positions; see caption, Figure 4. Results generated using all available data show the same trend, but data sets with many SNPs have tighter distributions.

**Figure 4:**
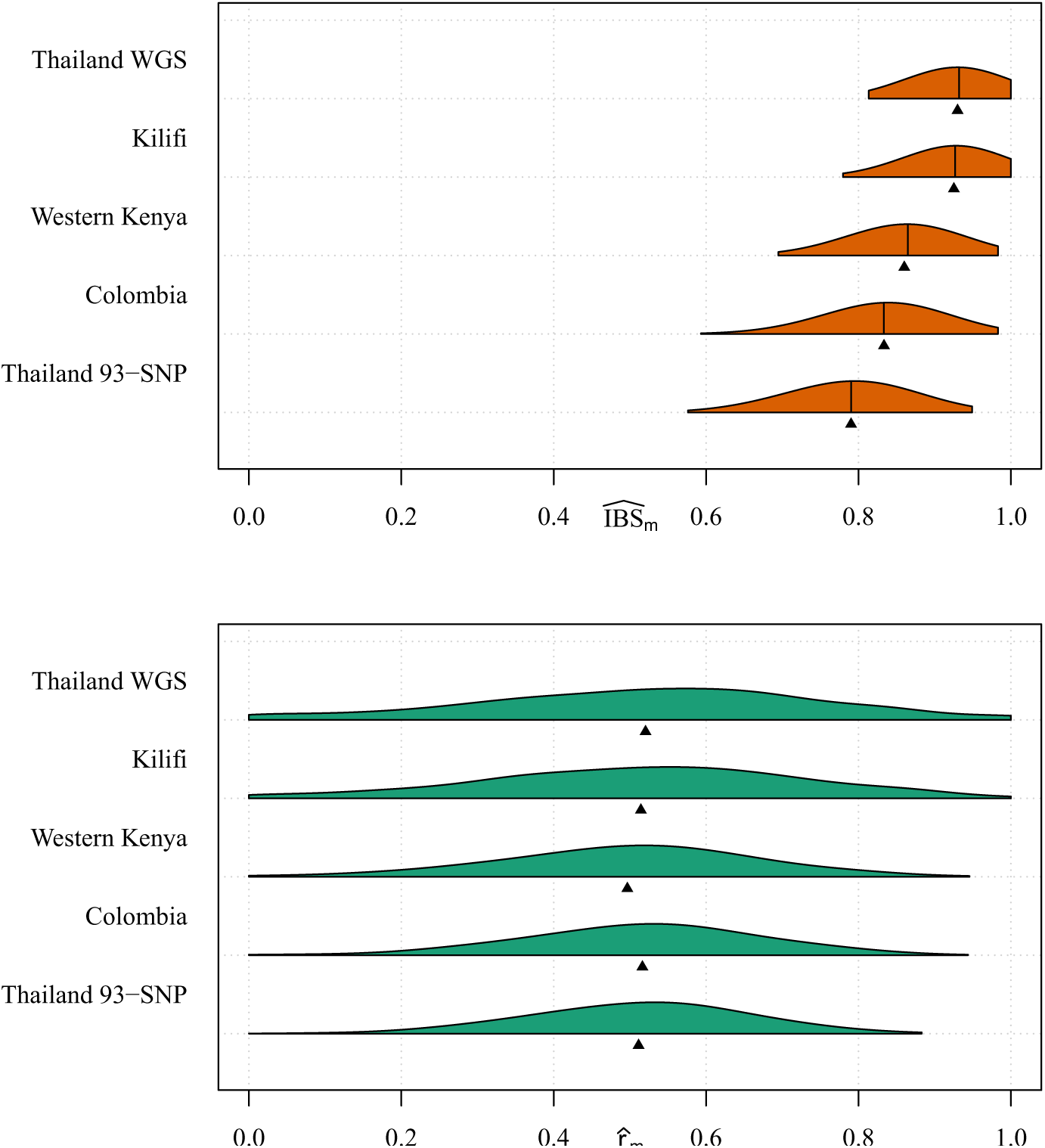
Measures of relatedness: pairs simulated with relatedness 0.5: Violin plots showing distributions of 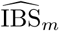 (top) and _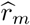_ (bottom) each based on 100 pairs simulated using *r* = 0.5 and allele frequency estimates based on *P. falciparum* data sets with at least 59 SNPs (Table 1). To single out the effect of frequencies, we fixed all other parameters across the data sets, including the number of SNPs simulated and their positions. Specifically, we used 59 SNPs whose positions were extracted from the Western Kenyan data set. Allele frequencies were sampled uniformly at random from the full set of allele frequency estimates based on each data set. For each set of 59-SNP allele frequencies, the 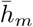 values were 0.86, 0.85, 0.73, 0.67, 0.58 (top to bottom row of each plot, respectively). Data were simulated under the HMM with *ε* = 0.001, *r* = 0.5 and *k* = 1. Black vertical bars denote 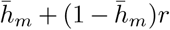 (top) and triangles denote the mean 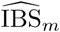 (top) and mean _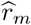_ (bottom).

Figure 5 shows 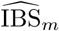 and 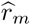 distributions based on the real sample pairs from published data sets. The location and spread of the 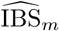 distributions vary considerably. As Figure 4 exemplified using simulated data, comparisons of absolute 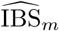 values are non-portable across data sets. It is thus wrong to interpret the left-most centering of the distribution based on the real 93-SNP data set from Thailand as evidence that *P. falciparum* parasites from Thailand are less related than those from Kenya, or that they represent a different population to that represented by the WGS data set also from Thailand. Despite very different absolute values of 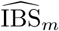, careful inspection shows all center around 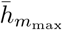, which is the expectation of 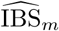 for unrelated parasites pairs whose *r* = 0 (equation 3.3). We thus conclude that many parasite pairs in these real data sets are unrelated. Our conclusion is corroborated by estimates of relatedness based on 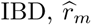 (Figure 5, bottom). Though the vast majority of parasite pairs are unrelated, we see some variation in the mean 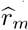. This variation is caused in part by outliers (parasite pairs with high relatedness). It is also caused by variation in 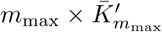: estimates of *r* are more error prone when 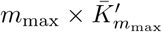 is small and those close to zero (the vast majority) are liable to upward bias due to boundary effects; see next section. The distribution of 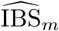 based on the *P. vivax* data set (Thailand MS) most closely approximates its partner distribution of 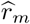 due to the highly polymorphic nature of the microsatellite data whose 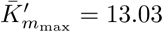.

**Figure 5:**
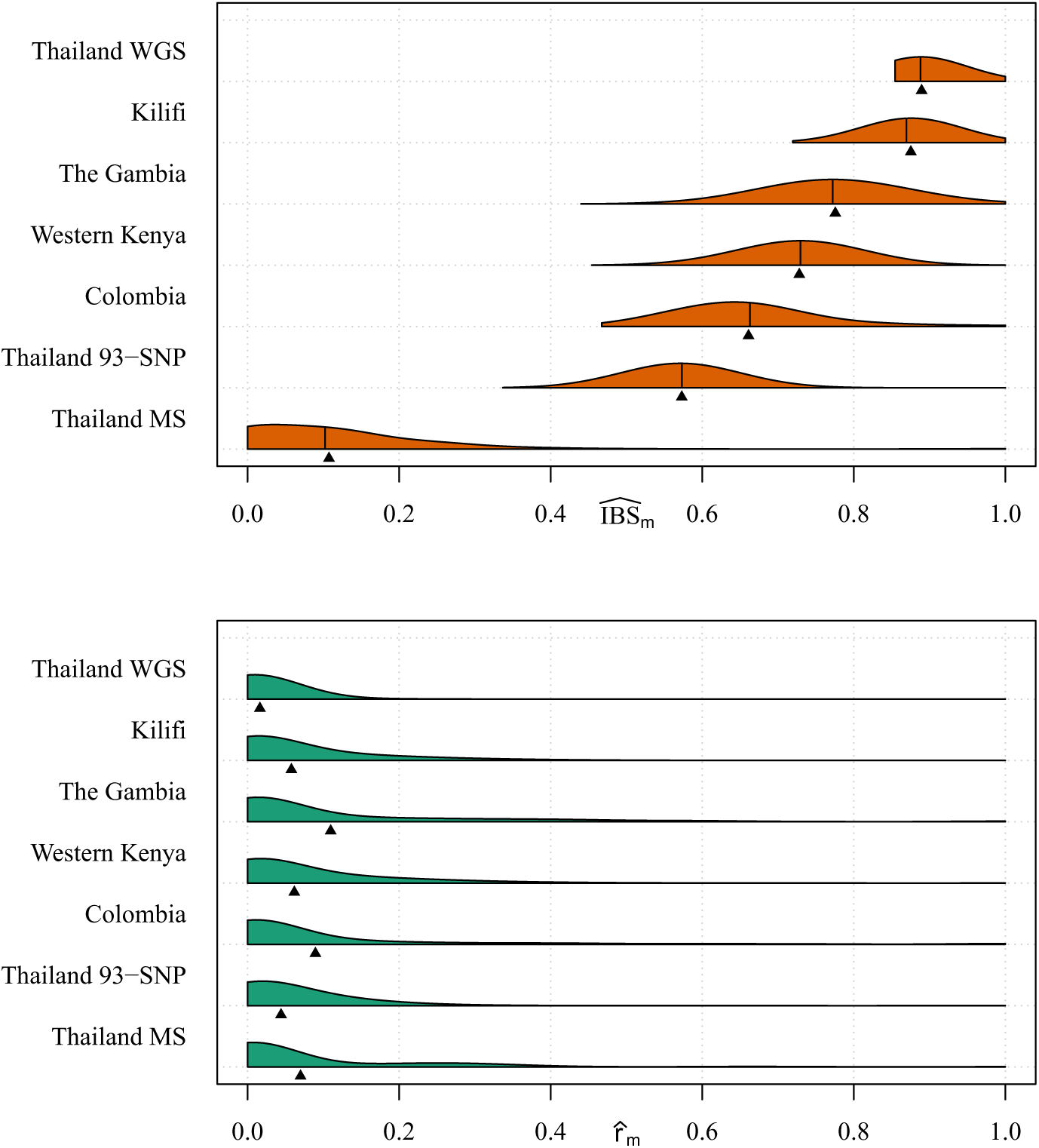
Measures of relatedness: parasite pairs with unknown relatedness. Violin plots showing distributions of 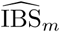 (top) and 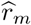 (bottom) based on pairwise comparisons of *Plasmodium* monoclonal samples from six published *P. falciparum* biallelic SNP data sets (Table 1) and a single *P. vivax* microsatellite data set (Thailand MS). Black vertical bars denote 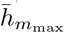(top) and triangles denote the mean 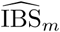 (top) and mean 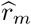 (bottom).

### 4.2 Estimating relatedness

Distributions of estimates of relatedness between pairs of *Plasmodium* monoclonal samples are plotted in Figure 5 (bottom plot). For each site, 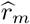 values range from 0 to 1, suggesting presence of unrelated, partially related and clonal parasites across all data sets. The vast majority, however, have 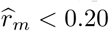. For a selection of 100 estimates ranging from 0 to 1, Figure 6 shows 95% parametric-bootstrap confidence intervals. In general, confidence intervals are tighter around estimates for data sets with larger 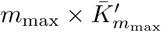, a point we shall return to later. Due to the asymmetric nature of confidence intervals near zero and one, estimates of *r* close to the boundary are liable to be biased. Considering the boundaries, intervals around estimates of *r* close to one are tighter, in general, than those for *r* close to zero.

**Figure 6:**
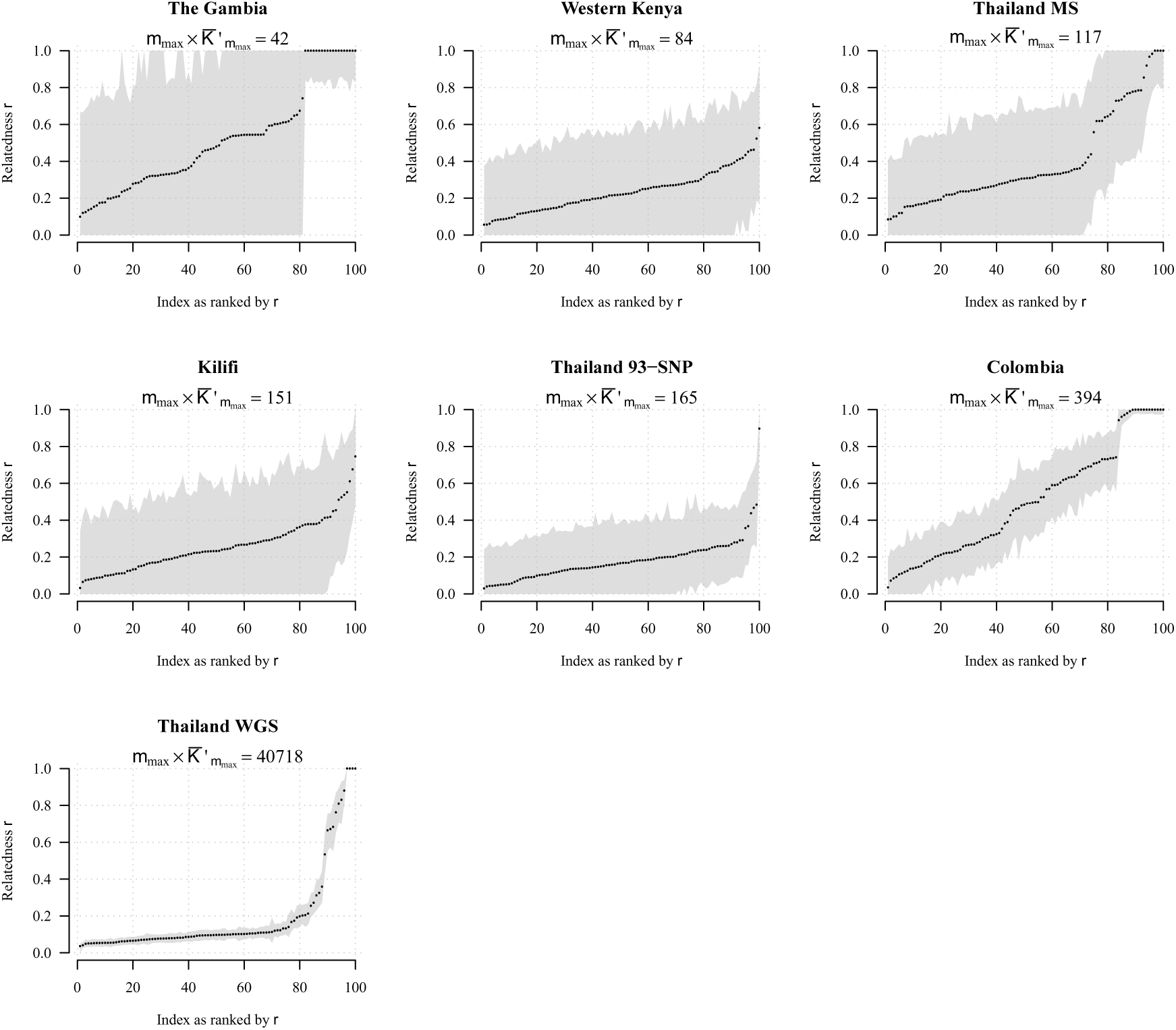
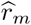 with 95% confidence intervals for 100 select pairwise comparisons of monoclonal *Plasmodium* samples from *P. falciparum* data sets list in Table 1 and a single *P. vivax* data set, Thai MS.

Considering Figures 5 and 6, we used the HMM to generate 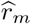 for biallelic marker data sets whose *m*_max_ is greater than 24 (Table 1), and the independence model to generate 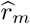 for the polyallelic data set whose *m*_max_ = 9, Thai MS. Based on simulated data, the HMM provides coverage^3^ close to 0.95 for *m >* 24, while the independence model provides waning coverage for *m >* 24, especially when *k*, which parameterizes the switching rate of the Markov chain, is small; for *m* = 24 both the HMM and the independence model provide similar coverage, above or around 0.85 (Figure 7).

**Figure 7:**
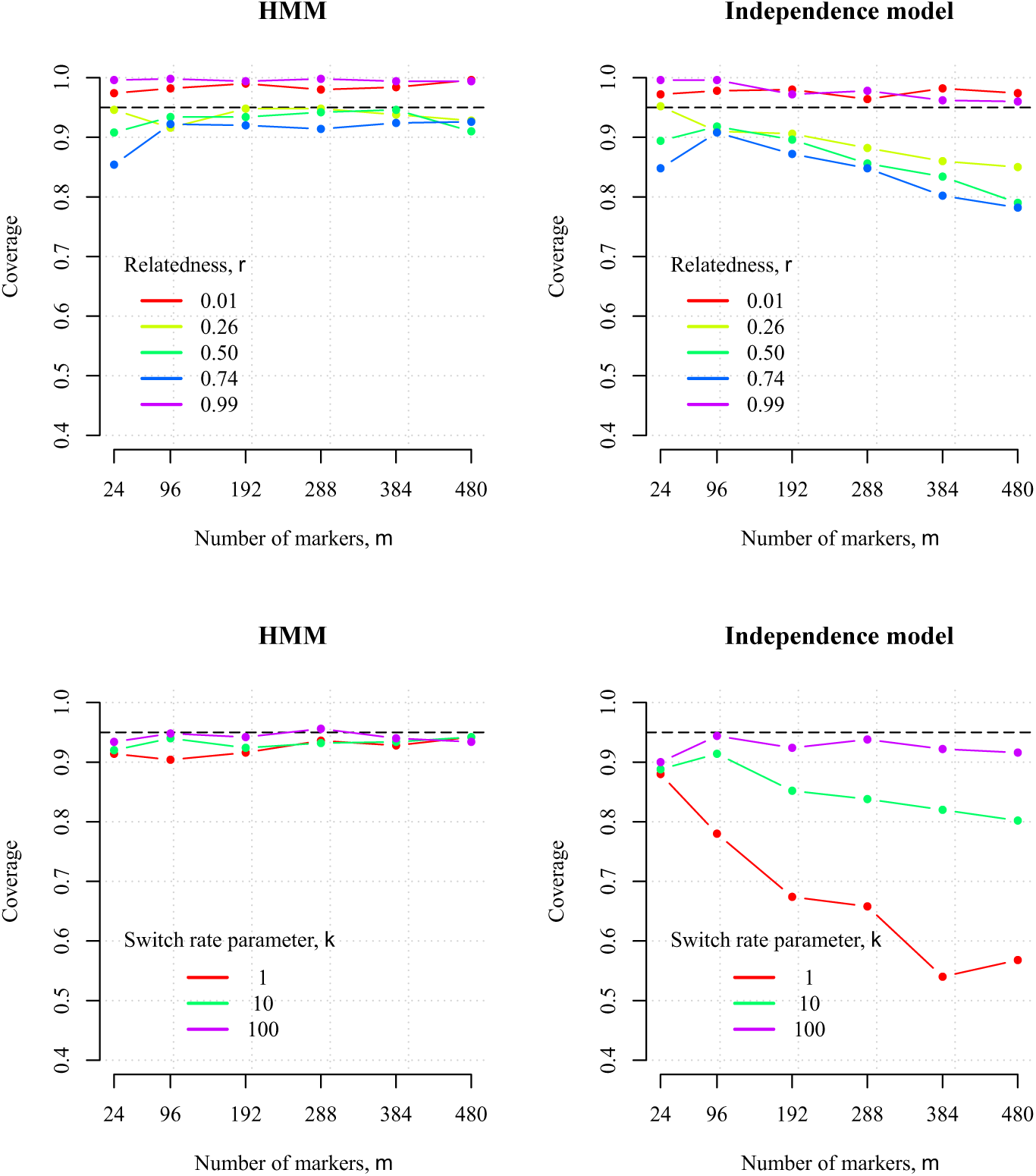
Coverage of 95% parametric bootstrap confidence intervals constructed under the HMM (left) and independence model (right). Coverage is equal to the proportion of 500 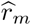 whose 95% parametric bootstrap confidence intervals contain the value of *r* used to simulate the data. It was based on data simulated under the HMM with *∈* = 0.001. Data were simulated for *m* biallelic markers (i.e. *K*_*t*_ = 2 *∀ t* = 1, …, *m*). Plots on the top show coverage for data simulated with different values of *r* given fixed *k* = 12. Plots on the bottom show coverage for data simulated with different values of *k* given fixed *r* = 0.5.

For all data sets biallelic and polyallelic, we construct confidence intervals using the parametric bootstrap due to the non-quadratic nature of the log-likelihood of *r* when 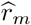 is close to either 0 or 1 (e.g. Figure B.3, left top and middle). For 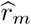 away from 0 and 1, the log-likelihood is quadratic (e.g. Figure B.3, bottom left plot) and thus normal-approximation confidence intervals could be constructed.

### 4.3 Marker requirements for prospective relatedness inference

As Figure 4 exemplified using simulated data, estimates of 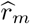 concentrate around the value of *r* used to simulate the data but with large variability, in part due to the finite length of the genome and in part due to limited data. We now consider how large *m* needs to be to estimate *r* with specified RMSE using different marker types (i.e. considering *K*_*t*_ = 2 *∀t* = 1, …, *m* and, more generally, *K*_*t*_ ≥ 2).

#### 4.3.1 Biallelic markers

First we consider relatedness inference using biallelic markers (e.g. SNPs, the most abundant polymorphic marker type, commonly used for relatedness inference [1]).

Figure 8 shows the RMSE of 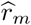 generated under the HMM given allele frequencies drawn with probability equal to their minor allele frequencies (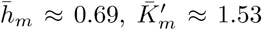) versus allele frequencies drawn uniformly at random (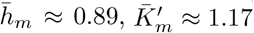). There are three notable results. First, errors obtained using allele frequencies drawn at uniformly at random are smaller (Figure 8, left). This is in agreement with the long-established result that higher minor allele frequencies are preferable for relationship inference [57]. Second, the RMSE is relatively large for 24 markers, decreasing dramatically upon increasing the marker count to 96. Though less dramatic, the decrease in RMSE is appreciable up to 288 markers, with diminishing returns thereafter. RMSE does not tend to zero due to the finite length of the genome. Third, RMSE error decreases with increasing proximity of the data-generating *r* to either 0 or 1 (especially the latter). As such, biallelic marker requirements for inference of *r* = 0.5 constrain guidelines for inference of *r* in general (Table 2).

**Table 2:** Biallelic marker requirements for specified RMSE around *r ∈* {0.01, 0.50, 0.99} and any *r ∈* (0, 1) extracted from Figure 8, left (i.e. given allele frequencies with 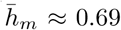. The length of the genome is denoted by *L*. *†*Since *r* = 0.5 has the largest marker requirements in general, inference of any *r ∈* (0, 1) is given by the maximum of the marker requirement interval for *r* = 0.5.

| RMSE | $r = 0.01$ | $r = 0.50$ | $r = 0.99$ | Any $r \in (0, 1)^\dagger$ |
| --- | --- | --- | --- | --- |
| 0.00 | $> L$ | $> L$ | $> L$ | $> L$ |
| 0.05 | 192-288 | $> 480$ | $< 24$ | $> 480$ |
| 0.10 | 24-96 | 96-192 | $< 24$ | 192 |
| 0.15 | 24-96 | 24-96 | $< 24$ | 96 |
| 0.20 | $< 24$ | 24-96 | $< 24$ | 96 |

**Figure 8:**
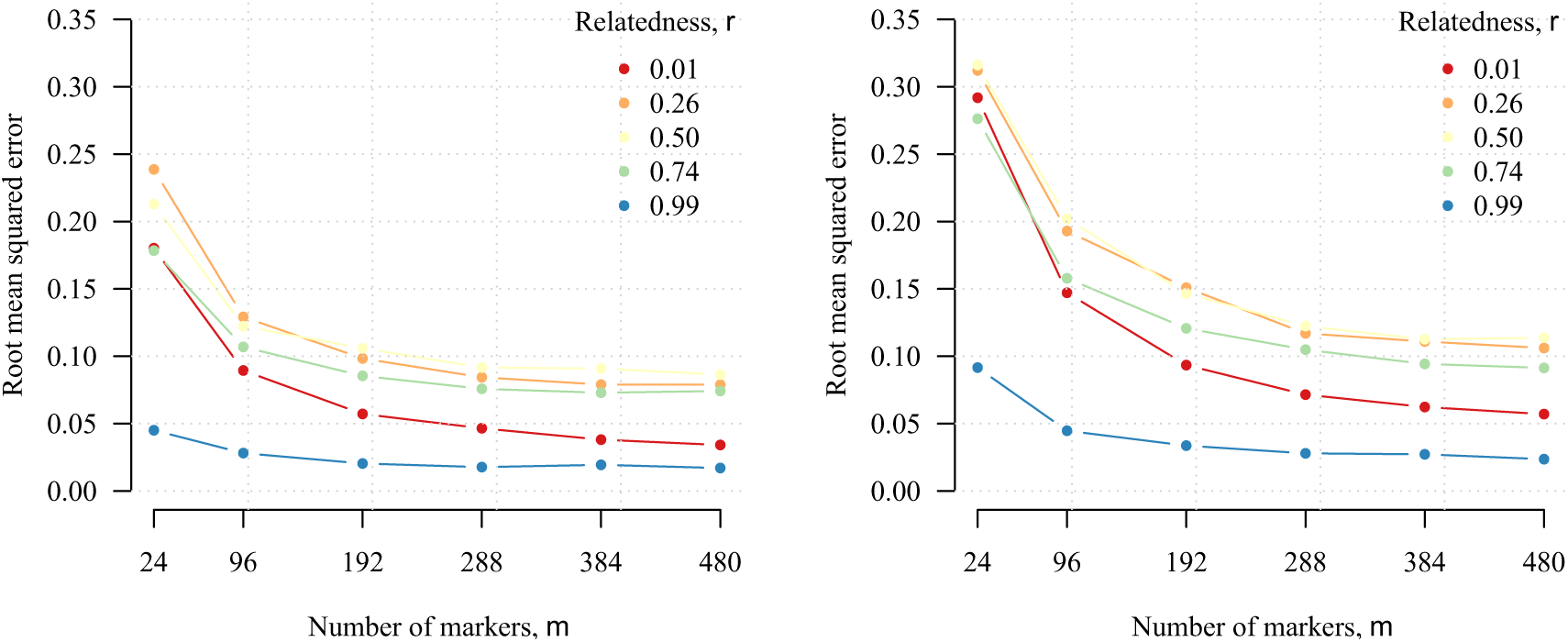
RMSE of 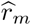 generated under the HMM. Data were simulated under the HMM using various *r* (see legend); allele frequencies with 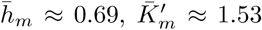 (left plot) and 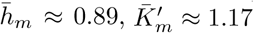 (right plot); *ε* = 0.001, *k* = 12, *K*_*t*_ = 2*∀t*.

#### 4.3.2 Polyallelic markers

Highly polyallelic microsatellite length polymorphism markers have long been used for relatedness inference, and there is growing interest in using microhaplotypes (short highly diverse regions of the genome) [1, 58]. Neither microsatellite nor microhaplotypes are point polymorphisms. However, to explore the general utility of polyallelic markers for relatedness inference, we make the simplifying assumption that they are.

Figure 9a shows three notable results. First, if only a small number of markers (e.g. 24) are available, a slight increase in their average effective cardinality markedly reduces RMSE, with diminishing returns as *m* grows. Second, to obtain RMSE less than some arbitrary amount, there may be an option between increasing *m* and increasing cardinality. For example, to obtain RMSE *<* 0.1, our results suggest typing 96 markers with 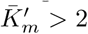 or around 192 markers with 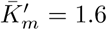 This latter option agrees roughly with the requirements for *r* = 0.5 in Table 2. Third, within the range of *m* values explored here, markers with *K*_*t*_ > 2 are necessary for optimally low RMSE (i.e. to achieve RMSE comparable with Mendelian sampling and thus negligible RMSE due to marker limitations).

**Figure 9:**
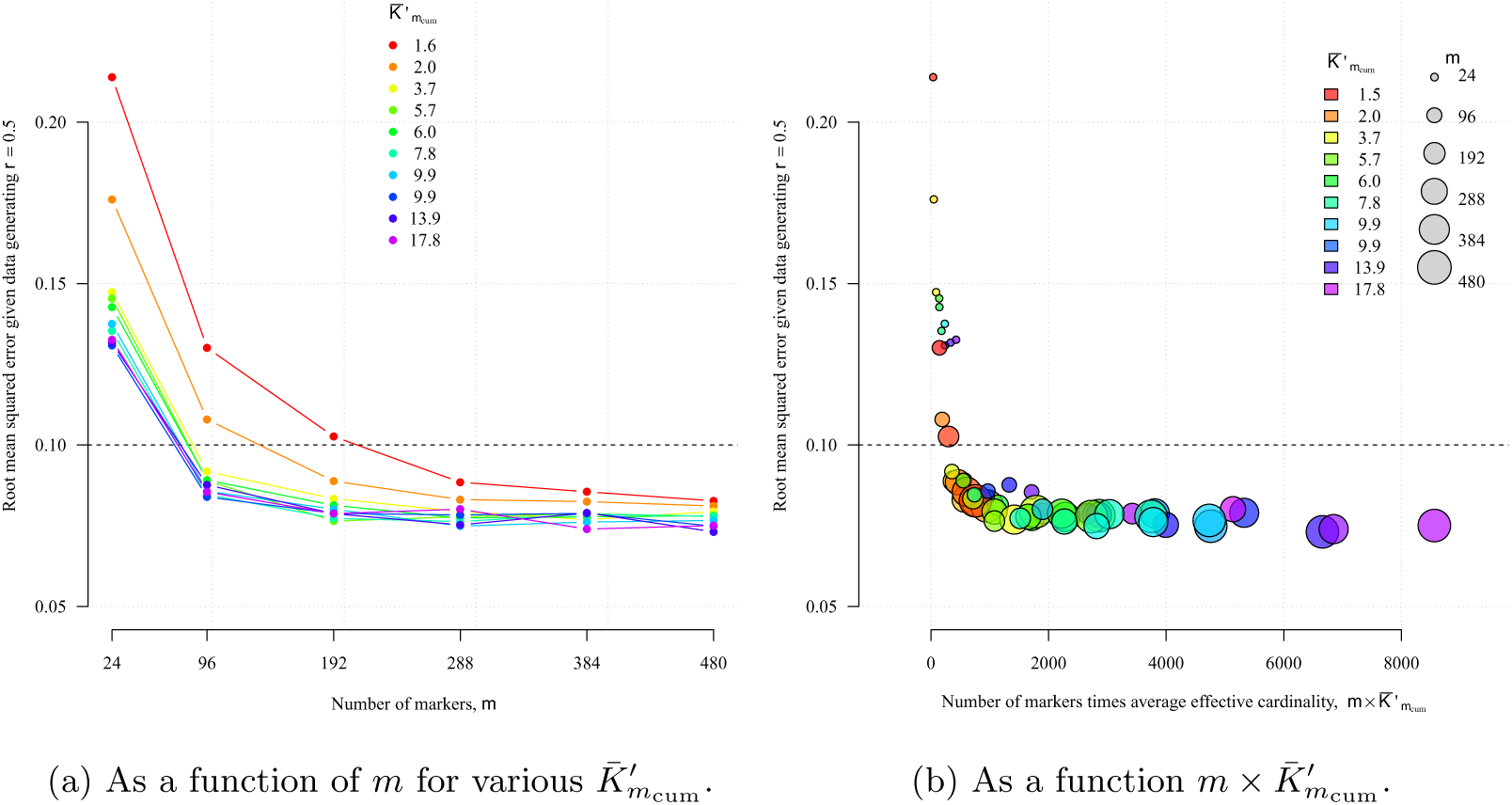
RMSE of 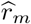 around data generating *r* = 0.5 with number of markers, *m*, and average effective cardinality,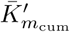.

The results shown in Figure 9a are projected onto a single axis in Figure 9b, showing the synergistic effect of increasing both *m* and 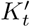. Clearly, larger 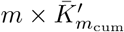 provides smaller RMSE with diminishing returns beyond 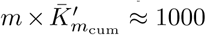. Informally, this result provides intuition as to why we obtain, in general, tighter confidence intervals around 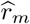 based on *Plasmodium* data sets with larger 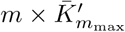(Figure 6). Moreover, it suggests that the confidence intervals around the Thailand WGS estimates are as small as they can be.

## 5. Discussion

Using a simple model framework, we call attention to properties of estimates of genetic relatedness, *r*, increasingly used in genetic epidemiology of malaria. These results, though articulated around monoclonal haploid malaria parasites, are applicable more generally to haploid eukaryotes (highly recombining prokaryotes would require a modified model).

The fraction IBS, which does not distinguish between alleles shared due to ancestry versus chance, is not a statistically principled estimator of *r*. As such, it does not allow calculation of confidence intervals for *r*, nor marker requirements. Its expectation is a correlate of *r*, but absolute values and quantitative estimates of trends are not portable across studies due to dependence on allele frequencies, which vary in space and time, and with different marker panels and quality control procedures [2]. On the contrary, measures based on IBS have the advantage of not depending upon potentially problematic allele frequency estimates discussed below [2, 4]. By illustrating how the fraction IBS is expected to change as a function of *r* and the alleles frequencies, we aid interpretation across studies using measures based on IBS to investigate relatedness.

Model-based relatedness inference allows construction of confidence intervals and marker requirements. Based on the parameters we explored, we recommend successful genotyping of at least 200 biallelic or 100 polyallelic markers for relatedness inference with RMSE less than 0.1 (if markers are highly polyallelic, fewer may be required, as in the Thai MS data set). In practice, a chosen set of makers could combine biallelic SNPs and more polyallelic marker types (e.g. microhaplotypes). Though not directly comparable, our results roughly agree (are of the same order of magnitude), with those reported for diploids and polyploids (Table 3). Relatedness inference for polyploids (e.g. [59, 13]) is comparable to that for polyclonal malaria samples, which arise due co-transmission and superinfection [60]. However, relatedness inference across polyclonal malaria samples is more challenging, since the equivalence of ploidy is unknown and variable. Despite these challenges, methods to infer relatedness within polyclonal malaria samples exist [23, 25], while methods to infer relatedness across polyclonal malaria samples are under development. It will be interesting to see how marker requirements, limited here to monoclonal malaria samples, scale in this more complex setting.

**Table 3:**
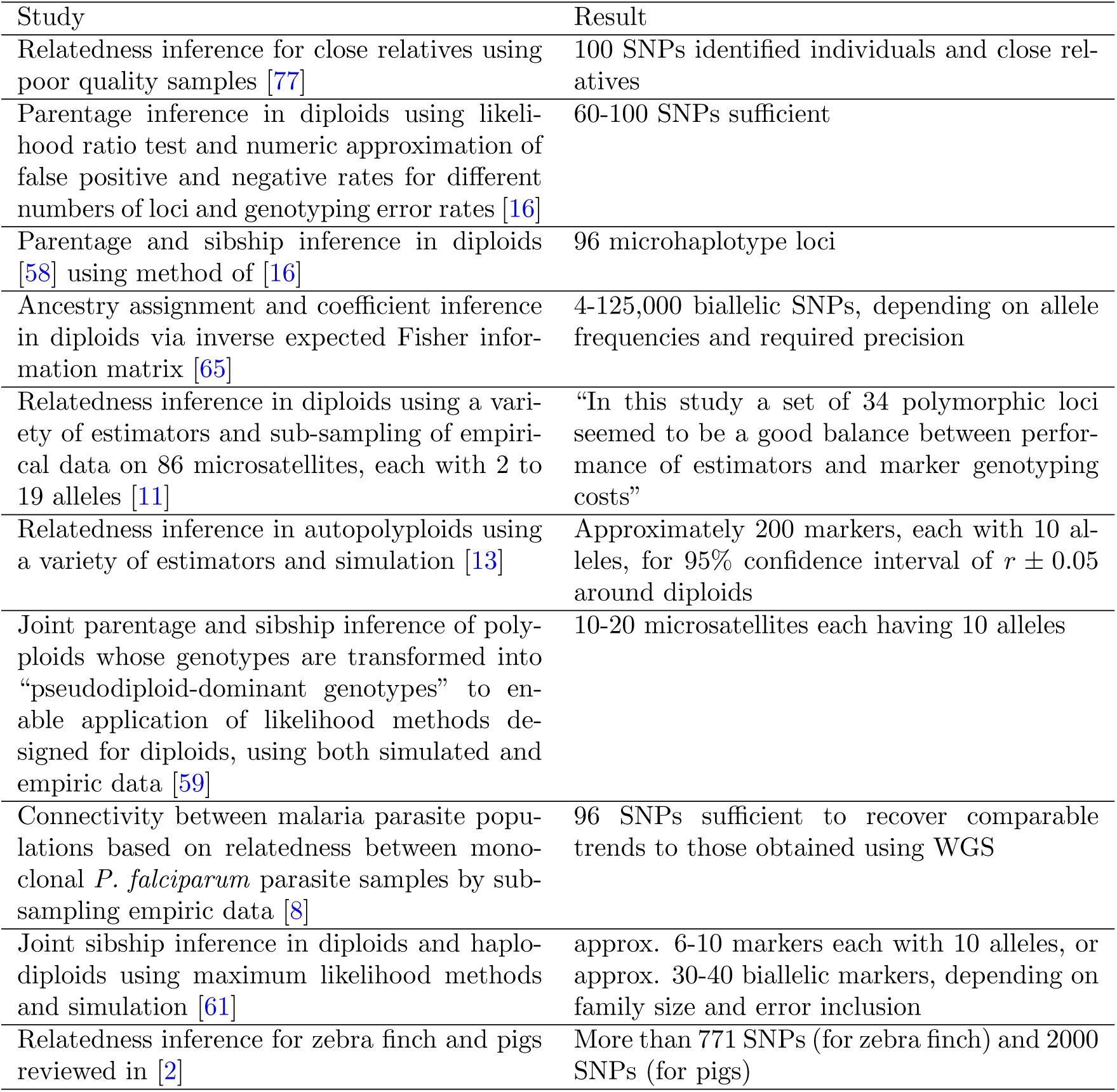
A non-exhaustive selection of studies in which numbers of loci for relatedness and associated inference are reported. Most of the above studies assume independence between markers because methods that assume dependence are, in general, designed for marker-rich applications where data requirements are not an issue.

| Study | Result |
| --- | --- |
| Relatedness inference for close relatives using poor quality samples [77] | 100 SNPs identified individuals and close relatives |
| Parentage inference in diploids using likelihood ratio test and numeric approximation of false positive and negative rates for different numbers of loci and genotyping error rates [16] | 60-100 SNPs sufficient |
| Parentage and sibship inference in diploids [58] using method of [16] | 96 microhaplotype loci |
| Ancestry assignment and coefficient inference in diploids via inverse expected Fisher information matrix [65] | 4-125,000 biallelic SNPs, depending on allele frequencies and required precision |
| Relatedness inference in diploids using a variety of estimators and sub-sampling of empirical data on 86 microsatellites, each with 2 to 19 alleles [11] | “In this study a set of 34 polymorphic loci seemed to be a good balance between performance of estimators and marker genotyping costs” |
| Relatedness inference in autopolyploids using a variety of estimators and simulation [13] | Approximately 200 markers, each with 10 alleles, for 95% confidence interval of $r \pm 0.05$ around diploids |
| Joint parentage and sibship inference of polyploids whose genotypes are transformed into “pseudodiploid-dominant genotypes” to enable application of likelihood methods designed for diploids, using both simulated and empiric data [59] | 10-20 microsatellites each having 10 alleles |
| Connectivity between malaria parasite populations based on relatedness between monoclonal *P. falciparum* parasite samples by sub-sampling empiric data [8] | 96 SNPs sufficient to recover comparable trends to those obtained using WGS |
| Joint sibship inference in diploids and haplodiploids using maximum likelihood methods and simulation [61] | approx. 6-10 markers each with 10 alleles, or approx. 30-40 biallelic markers, depending on family size and error inclusion |
| Relatedness inference for zebra finch and pigs reviewed in [2] | More than 771 SNPs (for zebra finch) and 2000 SNPs (for pigs) |

The results presented here are conditional upon an HMM under which various simplifying assumptions were made, the most significant being that of known and fixed allele frequencies. Typically, allele frequencies are estimated using data intended for relatedness inference yet assuming independent and identically distributed samples [61, 62]. These data-derived allele frequencies have been shown to give poor results and lead to underestimation of relatedness since “rare alleles shared by relatives are not recognized as such” [11]. Improving allele frequency estimates could benefit inference more than increasing the number of markers [11]. To better estimate allele frequencies of naturally occurring malaria parasites, for which pedigrees are unattainable, one could jointly model frequencies and relatedness as in [61]. Joint modelling would benefit inference in other ways also. For example, by borrowing information across samples and extending the inference framework, one could theoretically infer the ancestral recombination graph and thus the genetic map (presently assumed uniform across the malaria genome here and in [18, 23, 25]). That said, details specific to malaria (e.g. out-crossing versus selfing and their association with transmission) would present unique challenges (e.g. [63] and references therein). Modular extensions of pairwise methods to perform multi-way relatedness inference (e.g. [64]) have also been shown to outperform pairwise methods.

As formally stated in equation (3.6), we find that a highly polyallelic marker can be several times more informative than a biallelic marker for relatedness inference, comparable to results reported in population assignment [65]. Despite their superior informativeness, microsatellites are being superseded by SNPs due to the relative ease and reliability of typing the latter [1]. Recent interest in microhaplotypes (regions of high SNP diversity, unbroken by recombination) aims to combine the ease of SNPs with the informativeness of polyallelic markers [58]. Microhaplotypes can be defined *in silico*, using a decision theoretic criterion [66, 65], which relates to LD [48]. They can then be captured *in vitro* using amplicon sequencing [58, 67] or molecular inversion probes (MIPs), which can also be used to genotype microsatellites and SNPs [68, 69, 70]. The amplicon and MIP approaches are especially valuable for relatedness inference across multiclonal malaria samples, because amplicons and MIPs can capture within-host densities of different parasite clones as well as the phase of microhaplotypes in polyclonal infections [67, 70]. A model that accurately reflects the fact that microsatellites and microhaplotypes are not point polymorphism, while accounting for their associated mutation and observation error rates, thus merits consideration [71, 72].

Besides motif repeats within microsatellites and SNPs within microhaplotypes (presently overlooked), it is preferable to minimise dependence between markers. For any given *r* and *k*, dependence is a function of marker position and LD. As such, marker position is an important design consideration. When considering polyallelic markers, we sampled marker positions uniformly at random from the Thailand WGS data set. For microhaplotypes, a more realistic approach would draw from genomic intervals whose length is amenable to physical phasing and high LD. Doing so presents a trade-off between distance and window-wise effective cardinality. This trade-off is critical if diverse windows are genomically clustered. We do not consider it here, but it can be explored within the current framework and is the topic of future work. On the other hand, LD is a natural phenomena over which investigators have no degree of freedom. Some models commonly used in human genetics account for LD [46, 73] (also see [15]). Those designed to estimate relatedness between malaria parasites account for dependence between IBD states due to their physical proximity but not due to LD [18, 23, 25]. LD reported in malaria parasite populations (e.g. [74, 50, 75]) is generally lower than that reported in human populations [76]. Its incorporation into methods for malaria parasite relatedness inference, both within and between polyallelic markers, warrants further research.

Here and elsewhere marker requirements are based on either down-sampled or simulated data (Table 3). Standard asymptotic theory for HMMs is problematic in the present setting due to the finite length of the genome, and the increasing degree of dependencies between markers as their density grows. Understanding the finite sample properties of the maximum likelihood estimator in this setting remains an open problem. Another open problem beyond the scope of this study, is that of sampling individuals for population-level inference (e.g. how many parasite samples are required to reliably infer gene flow between different geographic locations using relatedness?). Work is ongoing to address these questions, which are very application-specific and dependent on many population factors (e.g. transmission intensity, seasonality, asymptomatic reservoir, etc.).

## 6. Conclusion

For portability, we recommend estimates of relatedness based on IBD for malaria epidemiology. To generate estimates between monoclonal parasite samples with less than 10% RMSE, approximately 200 biallelic markers or 100 polyallelic markers are required. Where studies inevitably differ in terms of available genetic data, confidence intervals illuminate inference. Together with anticipated work on population-level sampling, we hope this work on genetic-level sampling (and extensions thereof) will aid statistically informed design of prospective molecular epidemiological studies of malaria.

## 7. Acknowledgements

We thank all the authors of the *Plasmodium* data sets for either sharing their data or making them freely available online for use here and elsewhere. Pierre E. Jacob gratefully acknowledges support by the National Science Foundation through grant DMS-1712872. Aimee Taylor and Caroline Buckee are supported by a Maximizing Investigators’ Research Award for Early Stage Investigators, R35GM124715. This project was funded in part with Federal funds from the National Institute of Allergy and Infectious Diseases, National Institutes of Health, Department of Health and Human Services, under grant number U19AI110818 to the Broad Institute (Daniel Neafsey).

## Appendix A Estimator based on IBS

For clarity of exposition, here we derive results for 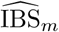 under a simple model that assumes no genotyping error (a more general result that includes genotyping error can be found in Appendix B, equation (B.9)). The simple model assumes the IBD state at the *t*-th locus, IBD_*t*_, is Bernoulli with relatedness parameter *r* ∈ [0, 1]. Given IBD_*t*_ = 0, we assume that 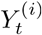 and 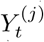 are independent Bernoulli with parameter 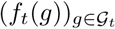. Given IBD_*t*_ = 1, we assume that 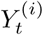 follows a Bernoulli with parameter 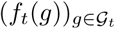 and that 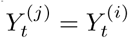 with probability one.

### A.1 Expectation of estimator based on IBS

In this section no assumptions are made about dependence between marker loci: equation (A.1) holds under both independence and dependence. The expectation of the estimator 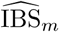 conditional on the frequencies 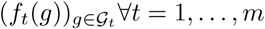 is

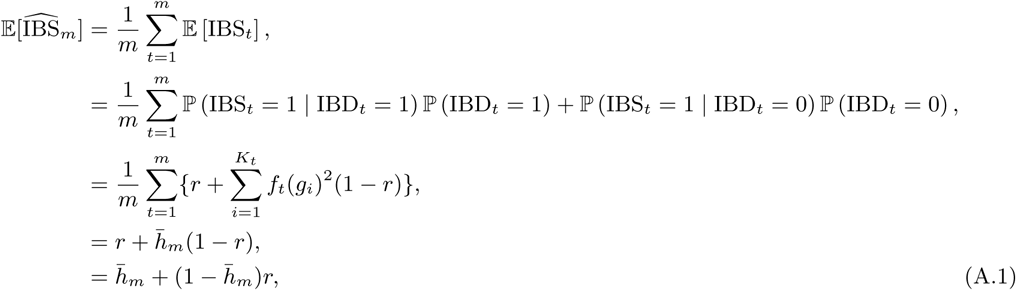

where 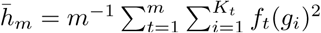 (equation (3.4)). Under different observation models, we would still obtain 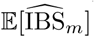 as a linear function of *r*; see second line above, where 𝕡 (IBS_*t*_ = 1 | IBD_*t*_ = 1) and 𝕡 (IBS_*t*_ = 1 | IBD_*t*_ = 0) could be anything as long as these expressions do not involve *r*.

### A.2 Convergence of estimator based on IBS

Here we work under the simplest setting: the measurements 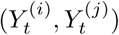 are independent across *t* = 1, …, *m*. In order to discuss convergence we need to imagine an asymptotic regime where *m* → ∞. We introduce an infinite sequence (*f*_*t*_(*g*_*i*_))_*t≥*1_, *i* = 1, …, *K*_*t*_, where each *f*_*t*_(*g*) is in (0, 1), and we introduce 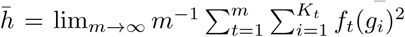, assuming the existence of that limit. To show that 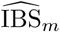 is not consistent for *r*, we show that it is consistent for 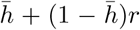, which is different to *r* unless *r* = 1. Thus we show that 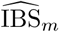 satisfies,

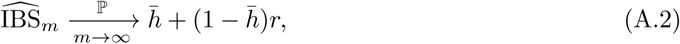

where the arrow is interpreted as “convergence in probability”. Since 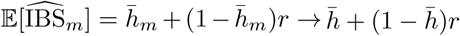 as *m* → ∞, we can establish (A.2) by showing that for every *ε* > 0

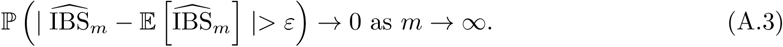

We show equation (A.3) by use of Hoeffding’s inequality (see Chapter 4 in [40]). Since 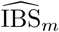 is an average of variables IBS_*t*_, which are bounded (IBS_*t*_ {0, ∈ 1}) and assumed independent, Hoeffding’s inequality yields

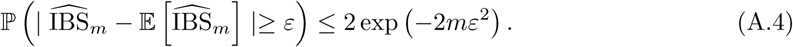

Since 2 exp (−2*mε*^2^) → 0 as *m* → ∞, equation (A.4) shows that equation (A.3) holds and therefore that equation (A.2) holds. Note that consistency could also be established in the dependent case, for instance via the application of a version of Hoeffding’s inequality for dependent processes.

Plots of 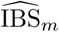 for data simulated under the independence model (Figure A.1) numerically show for *r* = 0 and 0.5 that 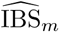 concentrates on its expectation (equation (A.1)) as more and more markers (*m* = 24, 96 and 192) are typed.

**Figure A.1:**
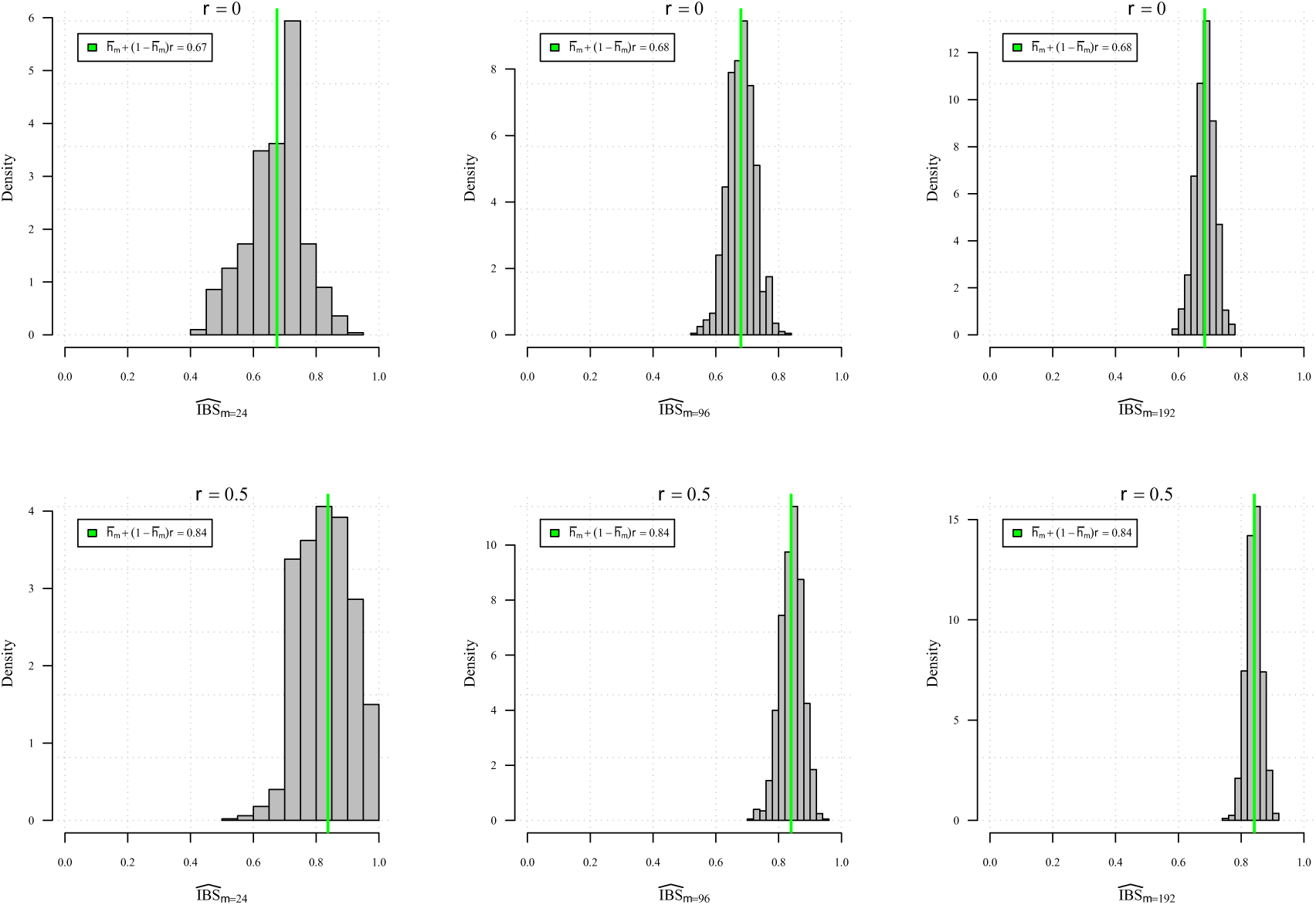
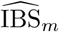 between pairs of biallelic marker data simulated under the independence model with different numbers of markers, *m*, and relatedness, *r*. The green vertical line marks 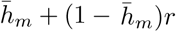 which is a function of the allele frequencies (equations (3.3) and (3.4)). Allele frequencies were sampled without replacement from Thai WGS data set with probability proportional to minor allele frequency estimates.

**Figure A.2:**
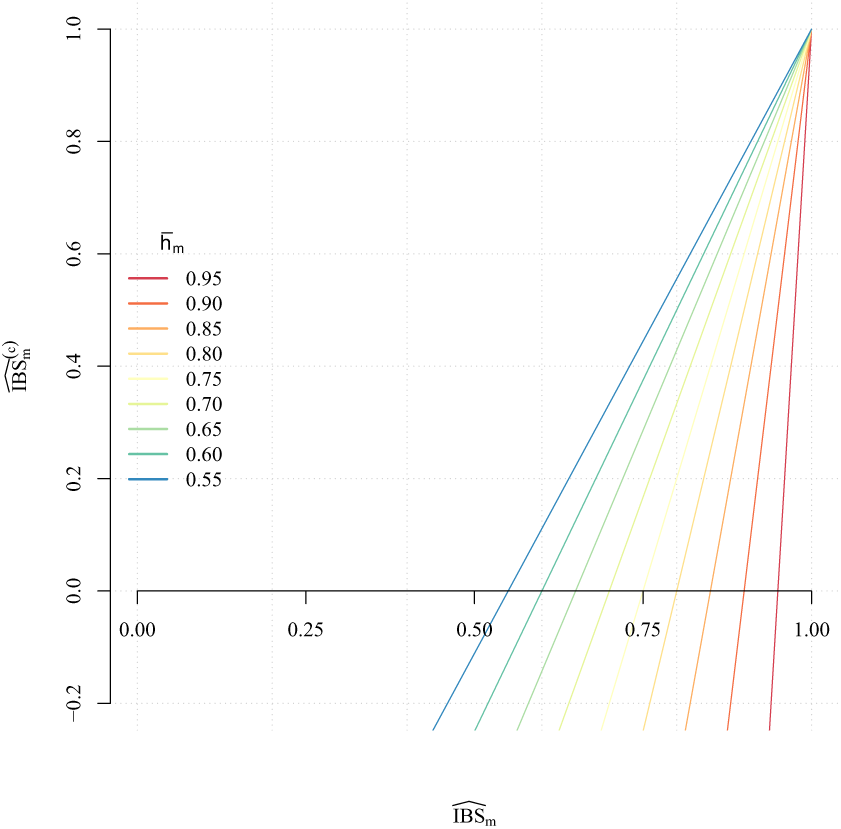
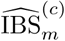 as a function of 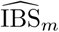 for various 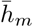 (equation (A.6)).

### A.3 Corrected estimator based on IBS

A corrected version of the estimator 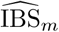 could be consistent for *r* (equation (A.7)) and is similar to existing method of moments estimators (reviewed in [11]), which generally underperform compared to maximum likelihood estimators (Chapter 9 of [40]).

By rearranging equation (A.1),

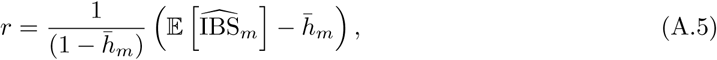

we can propose the following corrected estimator of *r*,

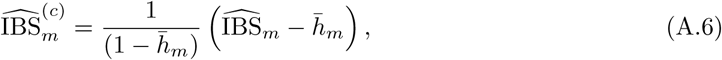

whose expectation is precisely *r*. The corrected estimator 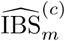 is consistent for *r*, with the same reasoning as in Appendix A.2 assuming independent observations,

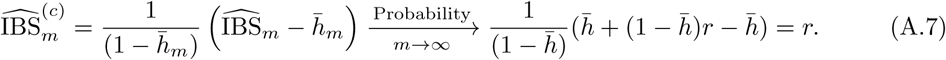

Figure A.2 shows a plot of equation (A.6) for different values of 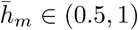. The range of 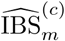 includes negative values. Setting negative estimates to zero can considerably improve results [11], but can also introduce bias [13]. For the *Plasmodium* data sets considered in the main text, Figure A.3 shows 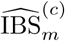 estimates truncated to [0,1].

**Figure A.3:**
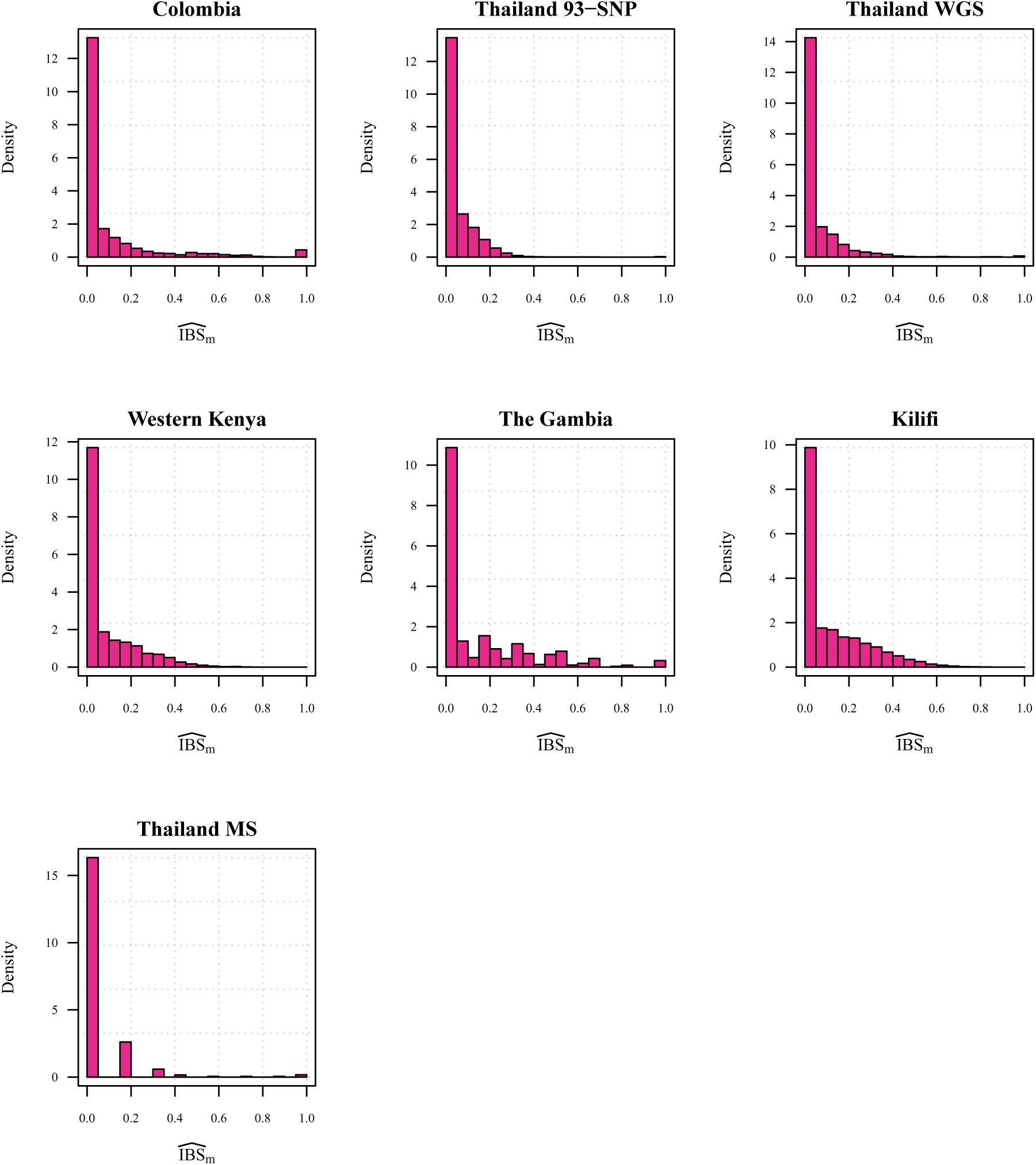
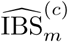 for several monoclonal *Plasmodium* data sets.

## Appendix B Model-based estimation of relatedness

### B.1 Framework

In this section we describe models that relate the available data to the objects of interest, in a self-contained presentation. The data comprise frequencies of alleles denoted by 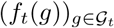, and allele indicators 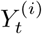, where the index *t* denotes a locus on the genome, and the superscript (*i*) refers to the *i*-th individual. The index *t* will run from 1 to *m*, the number of markers genotyped, and we will be particularly interested in the impact of *m* and *K*_*t*_ on the precision of the estimators. Note that *m* cannot be larger than *L*, the total length of the genome, which will create difficulties in making sense of an asymptotic regime where *m* goes to infinity, as will be discussed below.

We will consider pairs of individuals, *i* and *j*, for which we want to estimate the relatedness denoted by *r* and taking values in the interval [0, 1]. The models below might involve other parameters, and overall the vector of parameters is denoted by *θ*. We will make the first component of *θ* represent the relatedness *r*, so that *r* = *θ*_1_.

For each pair of individuals, we introduce a sequence of latent binary variables denoted by (IBD_*t*_) for identity-by-descent: IBD_*t*_ = 1 indicates identity-by-descent at locus *t*. We view this sequence as a two-state Markov chain. The case of independent variables for (IBD_*t*_) constitutes a particular case. In any case, the relatedness *r* ∈ [0, 1] represents the marginal probability that IBD_*t*_is equal to one, assumed to be identical for all *t*. While we do not observe (IBD_*t*_), we observe 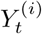 and 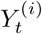 that are related to IBD_*t*_ at site *t* via an observation model, which can take into account the presence of genotyping errors. Together, the specification of the latent process (IBD_*t*_) and of the observation model fully describes a hidden Markov model, that can be used to estimate *r* using the data. Complete model specification is deferred to Appendix B.3, after a description of the general estimation procedure and some specific issues arising in the present case.

The estimation procedure is here based on the maximum likelihood approach. The likelihood function can be written as

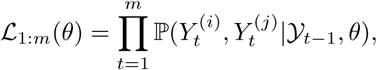

where 𝒴_*t-*1_ represents all the observations from locus 1 to locus *t* − 1, with the convention that 𝒴_0_is the empty set. We can further write each “incremental likelihood term” as

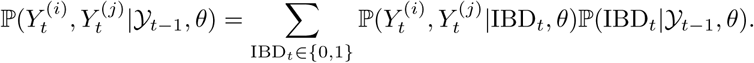

Since (IBD_*t*_) is a Markov chain, the forward algorithm [17] can be used to evaluate each incremental likelihood term for *t* = 1, …, *m*, for a cost of the order of *m* operations given *θ*.

We write *ℓ*_1::*m*_(*θ*) = log 𝓛_1:*m*_(*θ*), and 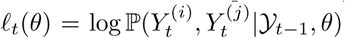. We denote the first and second derivatives of *ℓ*_*t*_(*θ*) by 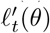 (a vector) and 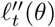 (a matrix) respectively. We will use the maximum likelihood estimator to approximate *r*, and we define it as

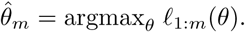

We next review some asymptotic properties of the maximum likelihood estimator (MLE) and detail how the present setting differs from the one usually considered in asymptotic studies.

### B.2 Distribution of the MLE

#### B.2.1 Standard asymptotic theory

We first recall what the usual asymptotic reasoning is for the distribution of the MLE in HMMs [35, 36, 37], in informal terms.

The first step is to imagine that the variables indexed by *t* (such as 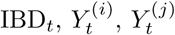, etc.) are part of infinite sequences of variables indexed by *t* ≥1. This allows us to consider a regime where the number of sites considered *m* can go to ∞. In Appendix B.2.2 we will discuss issues arising when applying this asymptotic reasoning in the present context of genetic data.

We observe that the log-likelihood and its derivatives are sums of *m* terms. Dividing by *m* yields averages, which might converge to limiting values as *m* grows large. For instance, the scaled log-likelihood might satisfy

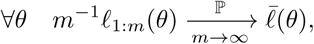

where the arrow is to be interpreted as “convergence in probability”, the left hand side of it being random if we consider the data to be random. Under some assumptions, the maximizer 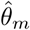 of *θ* ↦*m*^−1^*ℓ*_1:*m*_(*θ*) converges to the maximizer *θ*^*\**^ of the limiting function 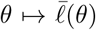. By the Taylor expansion of 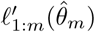 at *θ*^*\**^ we have

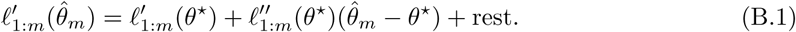

At the MLE 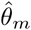, the derivative of the log-likelihood cancels: 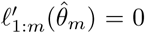, at least if the MLE is in the interior of the parameter space; extra care is required when the MLE is on the boundary of the parameter space, which occurs in the present setting where 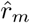 can be exactly zero or one. Therefore we obtain

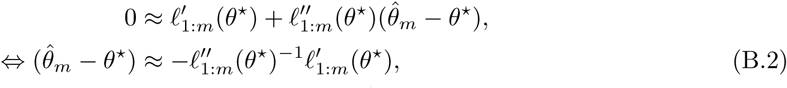

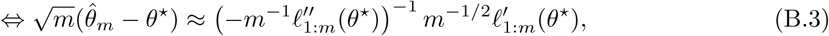

where ⇔ means “equivalently”. We will rely on the two following convergence results (see Chapter 13 in [78]),

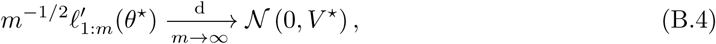

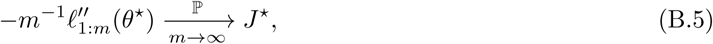

for some matrices *V* ^*\**^, *J* ^*\**^, assumed to be both semi-definite positive and symmetric. The first line above describes a convergence “in distribution” and can follow from a central limit theorem for the first derivative of the log-likelihood. The second line can follow from a law of large numbers applied to the second derivatives, as in Chapter 13 of [78]. We can combine these two convergence results using Slutsky’s lemma to obtain the asymptotic normality of the MLE:

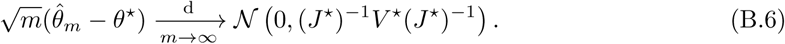

This key result can be used for sample size determination and for the construction of confidence intervals, provided that we can approximate *θ*^*\**^, *V* ^*\**^ and *J* ^*\**^ based on data. The asymptotic variance (*J* ^*\**^)^−1^*V* ^*\**^(*J* ^*\**^)^−1^ is sometimes called the sandwich formula, and can be estimated based on samples; see Doucet and Shephard [79] in the setting of hidden Markov models. If we assume that the model is well-specified, i.e. that the data actually are generated from the model with the parameter *θ*^*\**^, then it can be shown that *J* ^*\**^ = *V* ^*\**^ under regularity conditions (Chapter 13 of Douc et al. [78]). In this case, the asymptotic variance in (B.6) simplifies to (*J* ^*\**^)^−1^. The matrix *J* ^*\**^ is often termed the Fisher Information Matrix at *θ*^*\**^.

We briefly discuss the numerical obtention of 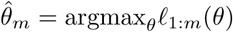. The log-likelihood function *θ* ↦ *ℓ*_1:*m*_(*θ*) can be plugged in a numerical optimizer, such as that implemented in the optim function of R. Evaluations of the log-likelihood function require runs of the forward algorithm on the data, for a cost of the order of *m* operations. Alternatively, one can also run an expectation-maximization algorithm, which involves calculating expectations with respect to the distribution of the latent process (IBD_*t*_) using the forward-backward algorithm [80], also called Baum-Welch in the context of HMMs [17]. If the parameter is small-dimensional, e.g. one or two-dimensional, a simple way of approximating the MLE consists in evaluating the likelihood (using the forward algorithm) on a grid of parameter values, and selecting the parameter associated with the highest likelihood.

The matrix *J* ^*\**^ can be estimated by 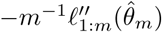, itself computed via numerical differentiation of the log-likelihood function at 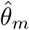. The estimation of *V* ^*\**^ is more complicated and has been the topic of a rich literature in time series analysis; see for instance Doucet and Shephard [79] and references therein.

#### B.2.2 Applicability of the standard asymptotic theory

The law of large numbers and central limit theorems usually employed to carry out the above reasoning, i.e. to establish (B.4) and (B.5) leading to the asymptotic normality of the MLE in (B.6), might not be meaningful in the present context. Indeed they usually apply to stationary processes observed over increasingly long periods of time. In such asymptotic setting, one eventually observes a realization of a stationary stochastic process over an infinitely long time horizon, which is enough to learn the invariant distribution of the process. We refer to this setting as standard asymptotics. Recall that our primary object of interest is the parameter *r*, which characterizes indeed the invariant distribution of the Markov chain (IBD_*t*_).

In the present setting where data comprise genetic sequences, increasing *m* means considering more loci on the genome. The *m* considered loci are located within the genome whose length is, however, fixed. Therefore increasing *m* amounts to increasing the subsampling frequency at which data are observed. In other words it decreases the distance between successive observed loci. We refer to this as subsampling asymptotics. To see where this differs from standard asymptotics, consider a simpler context where (IBD_*t*_) would not be hidden but directly observed. In the limit *m*→ ∞ in subsampling asymptotics, we would observe a continuous trajectory of (IBD_*t*_), switching from state 0 to state 1 and back again, over a fixed interval. The maximum likelihood estimate of *r* for such a model would be the proportion of time that the trajectory would spend in state 1 [81]. However this would not be exactly equal to *r*, even if the trajectory was sampled from the Markov model given *r*, because the fully-observed realization of (IBD_*t*_) would still be of a finite length; this is well-known, see [34] on the impact of the genome length on relatedness estimates under Mendelian sampling. On the other hand, in the standard asymptotics *m* → ∞ we would observe an infinitely long trajectory of the Markov chain, for which the maximum likelihood estimator of the transition matrix is consistent. The difference between the two regimes is illustrated in Figure

**Figure B.1:**
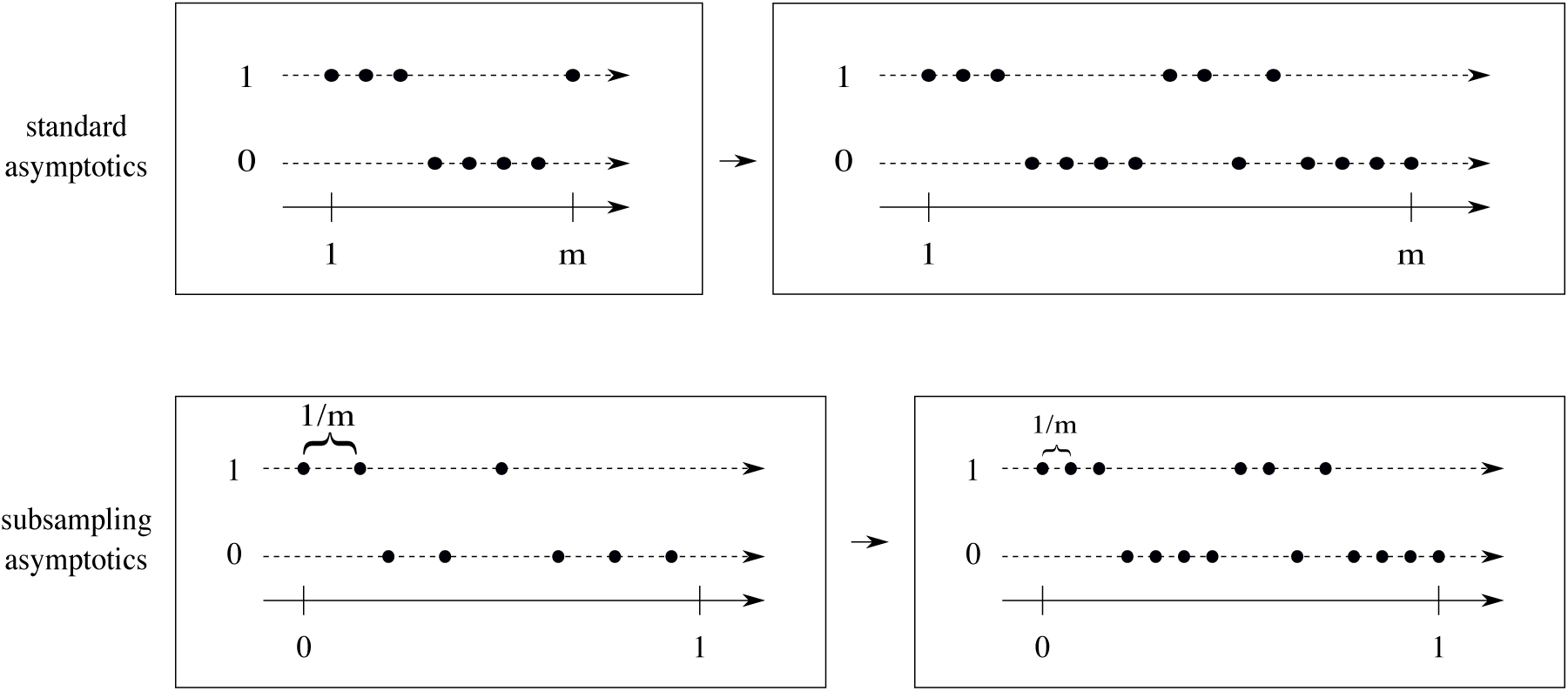
Two different ways of increasing *m*: in the top row, *m* refers to the length of the observation period, while the observations are separated by one unit of time. In the bottom row, the length of the observation interval is fixed to one, and the observations are placed at distance 1*/m* of one another; thus an increase in *m* means that successive observations are closer to one another, but the length of the observation period is fixed.

The difference in asymptotic regimes has consequences on the estimability of *r*. In the subsampling asymptotics, it is impossible to arbitrarily decrease the error of 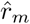 by increasing *m*: there is only so much information that can be gathered about *r* by increasing the number of loci under consideration; hence the distinction between expected IBD and realised IBD in [2]. A result such as the asymptotic normality with a 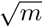 rate of convergence, as in (B.6), is in fact unlikely to hold. The numerical experiments indeed suggest that the root mean squared error associated with 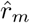 does not decrease beyond a certain point, no matter how large *m* is. The subsampling asymptotic regime has been formally studied with various applications to financial econometrics [82, 83], but we are not aware of similar results for hidden Markov models such as the ones considered here.

Despite the standard asymptotic results not holding, we do observe that the distribution of 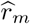 is approximately normal for *m* large enough (Figure B.2). This can be partially explained by the fact that normality of the MLE depends entirely on the log-likelihood being approximately quadratic [38], which itself does not have to follow from standard asymptotic arguments. Since the log-likelihood function is observed to be approximately quadratic providing 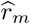 is not close to the boundaries (Figure B.3), we can still quantify the precision of the MLE by considering the second derivative of the log-likelihood at its maximum. Thus we will rely on the Fisher Information Matrix as a proxy for the precision of the MLE, in particular for the study of the effect of *K*_*t*_ in Appendix B.3.4.

**Figure B.2:**
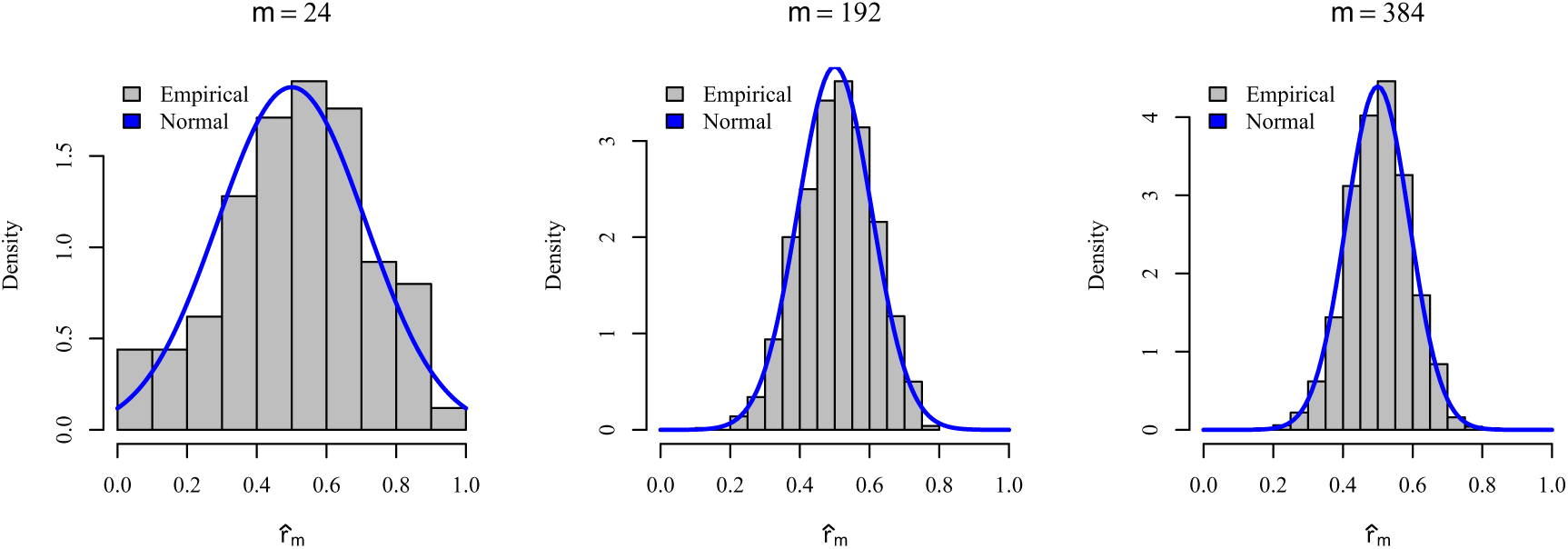
Empirical distributions of 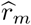 for different numbers of markers, *m*. Each distribution is based on 1000 estimates of *r* given data simulated and analyzed under the HMM with *r* = 0.5, *k* = 12, *K*_*t*_ = 2 *∀t* = 1 …, *m* and *ε* = 0.001.

### B.3 Models

We now describe a Markov chain model for (IBD_*t*_), followed by observation models for 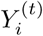 and 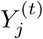 given IBD_*t*_.

#### B.3.1 Hidden Markov model

We write the transition probabilities of (IBD_*t*_) at a locus *t*,

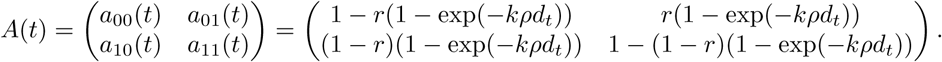

In the above, *a*_*jℓ*_(*t*) refers to the probability of IBD_*t*_ = *ℓ* given that IBD_*t-*1_ = *j*.

In the above expression, the relatedness is denoted by *r*; *d*_*t*_ denotes a genetic distance in base pairs (bp) between sites *t* −1 and *t*; *k* > 0 parametrizes the switching rate of the Markov chain and *ρ* is the recombination rate, assumed known and fixed across both haploid genotypes with value 7.4 *×* 10^−7^M bp^−1^ for *P. falciparum* parasites [33].

We can check that, if ℙ (IBD_*t-*1_ = 1) = *r*, then

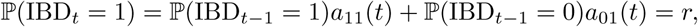

and thus the invariant marginal distribution of the chain is given by ℙ (IBD_*t*_ = 1) = *r*.

The above transition probabilities are at the core of many HMMs of relatedness (e.g. [14], where *k × ρ* = *a* and genetic distance *d*_*t*_ = *t*_*k*_ is measured in centi Morgans (cM), plus many subsequent models (see [15]), including [18], where *r* = *π*_1_ and 1 – *r* = *π*_2_.

We can check that, as the distance increases to infinity, the probabilities in *A*(*t*) simplify and correspond to the i.i.d. Bernoulli model where IBD_*t*_ is equal to one with probability *r*, independently for each site *t*. In other words, if sites are distant enough, we expect the HMM and the independence models to give similar results. This will happen in particular when *m* is small and when the loci under consideration are well-spread across the genome.

**Figure B.3:**
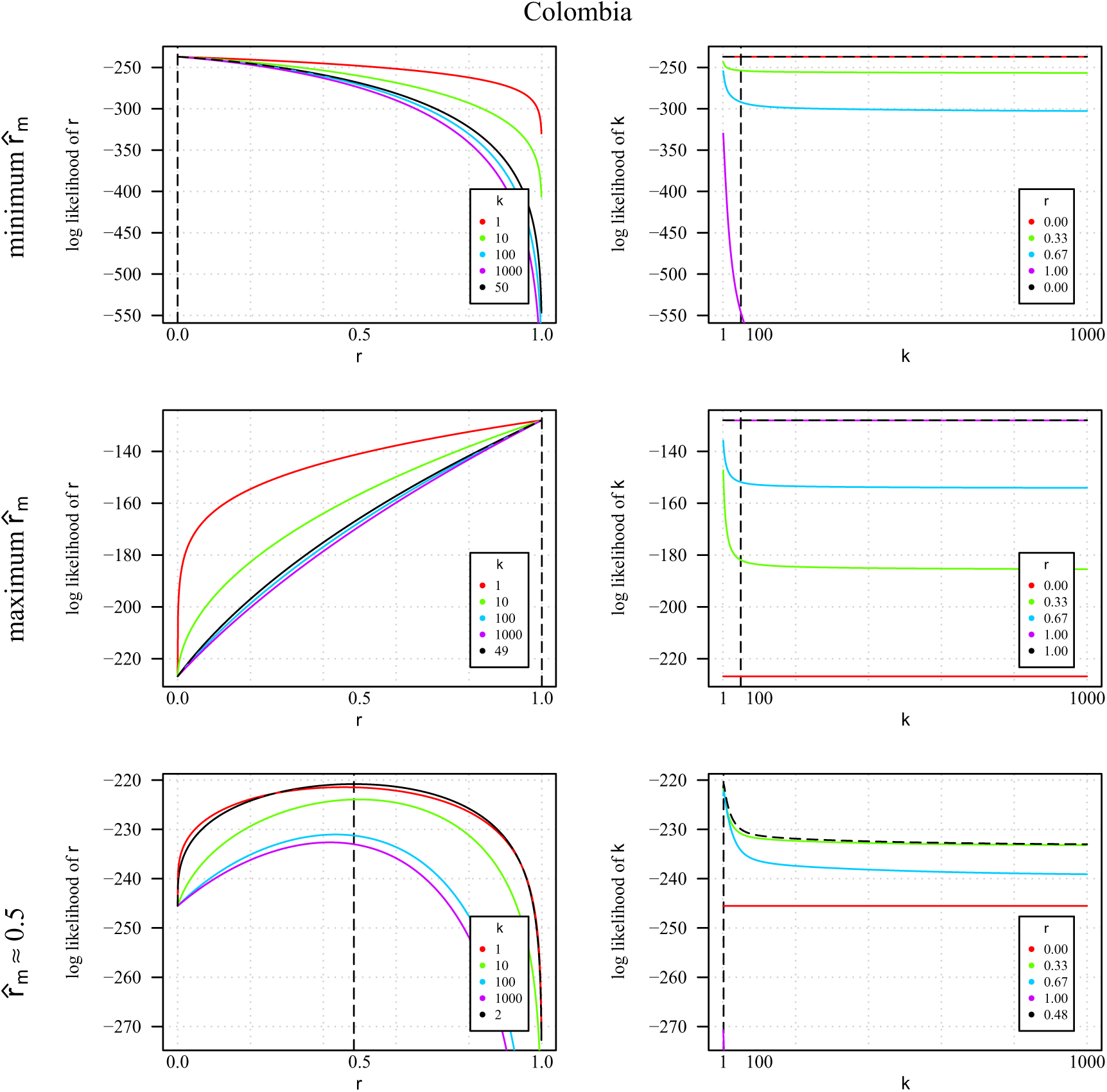
The log likelihoods of *r* for different *k* (left column) and *k* for different *r* (right column) for three different example sample pairs from the the Colombian data set: a sample pair with minimum 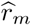 (top row, *m* = 248), a sample pair with maximum 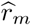 (middle row, *m* = 246), and a sample pair with 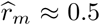 (bottom row, *m* = 245). Differences in *m* are due to missing genotype calls in the data. Vertical black dashed lines mark 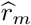 (left column) and 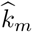 (right column). Black dashed function lines show the log likelihood of 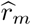 given 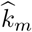 (left column) and of 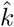 given 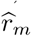 (right column). Coloured function lines show the log likelihood of 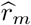 given values of 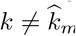 (left column) and of 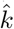 given values of 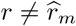 (right column).

#### B.3.2 Observation model

The observations 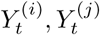 are related to (IBD_*t*_) only through IBD_*t*_ at locus *t*. The observation model introduces some true genotypes 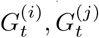 given IBD_*t*_, and then some genotyping error model defining the distribution of 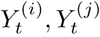 given 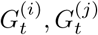.

First, the variables 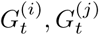 given IBD_*t*_ are defined as follows. If IBD_*t*_ = 0, then 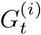 is inde-pendent of 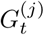 and both follow a Categorical distribution: for a set of values 𝒢 = {*g*(1), …, *g*(*K*_*t*_)} and probabilities {*f*_*t*_(*g*)} for *g* ∈𝒢, we have 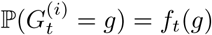, and likewise for 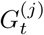. If there are only two types (e.g. the case for biallelic SNPs) then it is a Bernoulli distribution. If IBD_*t*_ = 1, then 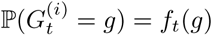 and 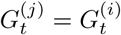 with probability one. Overall we can write the model as

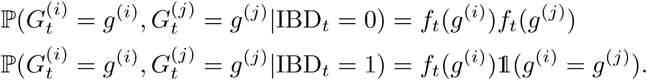

Next, we assume that genotyping errors occur independently for both individuals. This differs to the typical ‘all-or-none’ diploid setting (e.g. [14, 15]), since haploid genotypes in monoclonal parasite samples are genotyped separately. If they occur, we do not observe 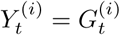 but instead we observe another genotype taken uniformly among the other possible values (by assumption); and likewise for the other individual *j*. This can be written

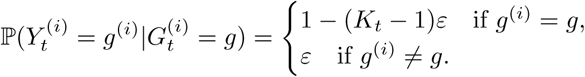

In the above expression *K*_*t*_ refers to the number of possibilities, which could be different for different sites *t*, and *ε* refers to a parameter such that the error rate is (*K*_*t*_ − 1)*ε*. This is suited to microsatellites in the sense that the error rate scales with *K*_*t*_ [72]. For biallelic SNPs, it amounts to a simple miscall.

Overall we can thus think of the observation model as the combination of a model for 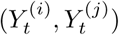 given 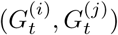 and a model for 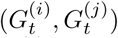 given IBD_*t*_. We can integrate 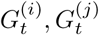 out to obtain directly the probabilities of 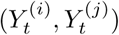 given IBD_*t*_:

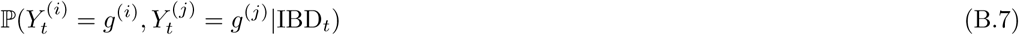

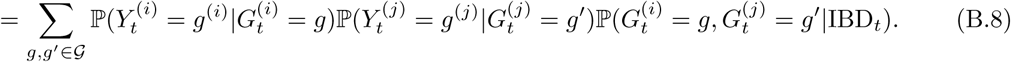

The cost of evaluating this expression is quadratic in the cardinality of. 𝒢

This observation model is the same (besides notation) as that for within-population samples under the HMM of hmmIBD [18] and, if *K*_*t*_ = 2, the same as that of the HMMs of isoRelate [23]. Mutations do not feature in it. However, any that do occur can be absorbed as errors, as they are considered to be in [61]. That said, it does not take into account microsatellite mutations in the sense that they scale with both motif size and repeat number [71], nor the inherent ordinal nature of microsatellites or the bias with regards to their amplification [84]. Bespoke adaptations could be made for specific data types.

### Digression: expection of fraction IBS considering error

Equation (B.8) means that

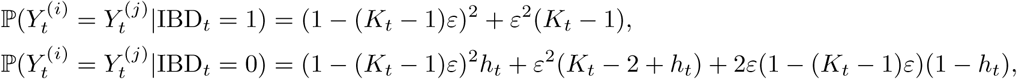

where 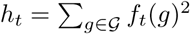. Consequently, under the present observation model,

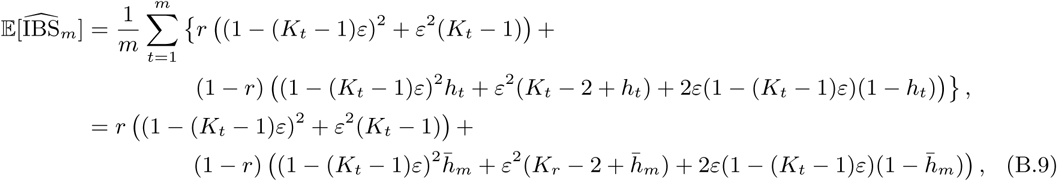

where 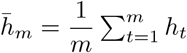 Equation (B.9) reduces to (A.1) when *ε* = 0.

#### B.3.3 The likelihood under the independence model

This model assumes independent random variables IBD_*t*_ across loci *t* ∈ {1, …, *m*}. It is a particular case of the above HMM when all *d*_*t*_ = ∞. Given a relatedness parameter *r* ∈ [0, 1], IBD_*t*_ is assumed Bernoulli with parameter *r*. Next, we define an observation model: given IBD_*t*_ = 0, we assume that 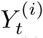 and 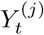 are independent Bernoulli with parameter *f*_*t*_(*g*). Given IBD_*t*_ = 1, we assume that 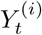 follows a Bernoulli with parameter *f*_*t*_(*g*) and that 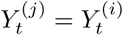 with probability one. This defines the observation model. The associated likelihood at site *t* is

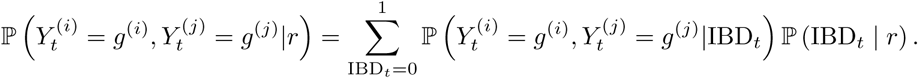

At this point we can define, for all *t*,

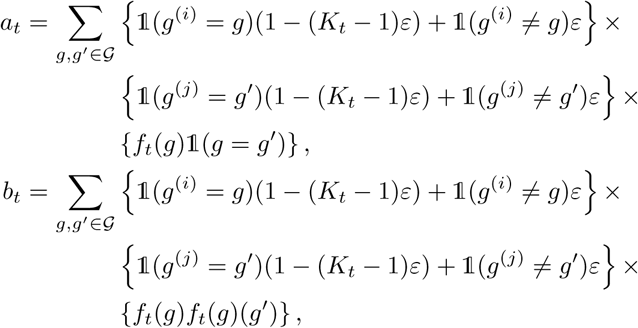

so that the likelihood reads 𝓛_*t*_(*r*) = *a*_*t*_*r* + *b*_*t*_(1 - *r*). The full log-likelihood can be simply written as

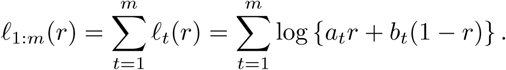

The gradient of the log-likelihood looks like

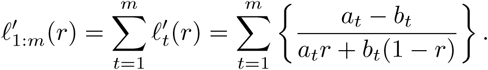

The second-order derivative of the log-likelihood looks like

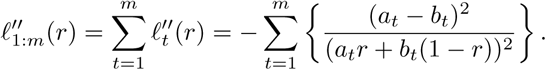

Since both numerator and denominator of each term are positive, 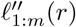 is strictly negative for all *r* ∈ (0, 1), and thus the function *r* ↦*ℓ*_1:*m*_(*r*) is concave on (0, 1).

For the HMM model, the form of the likelihood is less explicit; we do not have an explicit formula giving the likelihood as a function of *r* and of the data. However this is not a real problem as we can still numerically evaluate the likelihood, using what is usually called the forward algorithm [17]. Being able to numerically evaluate the likelihood leads to being able to optimize it to get the MLE, and to numerically differentiate it as well.

#### B.3.4 Maximizing Fisher information

We focus on a single site *t*, which we suppress from the notation. Let us denote the log-likelihood by *ℓ* and recall the formula

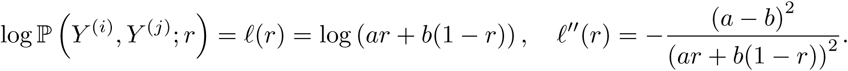

Assume there is no genotyping error for simplicity. Then *a* = *f* (*Y* ^(*i*)^) 𝟙 (*Y* ^(*i*)^ = *Y* ^(*j*)^) and *b* = *f* (*Y* ^(*i*)^)*f* (*Y* ^(*j*)^). From there the Fisher Information Matrix (FIM) is obtained as

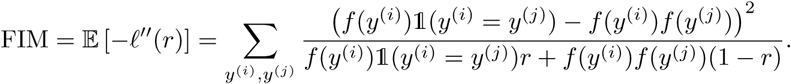

It is a function of *r* and of the allele frequencies. The FIM is proportional to the inverse of the asymptotic variance of the MLE, thus if we want precise estimators of *r*, we want a large FIM. This leads to the idea of maximizing FIM with respect to *f* for all *r*, to see which allele frequencies allow the best estimation of *r*. We can split the sum into the case for which *y*^(*i*)^ = *y*^(*j*)^ and the case for which *y*^(*i*)^ ≠*y*^(*j*)^; for simplicity we denote *f* (*y*^(*i*)^) by *f*_*i*_, which leads to

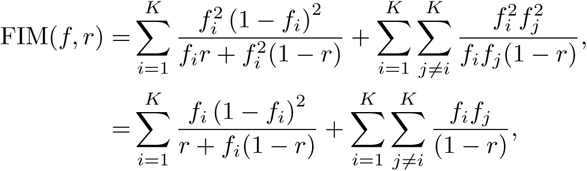

where we recall that *K* denotes the number of possible alleles. We note that Σ_*j*≠*i*_*f*_*j*_=1-*f*_*i*_ because 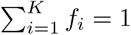, therefore we obtain

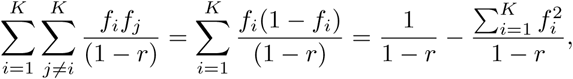

and thus the simpler form for the FIM:

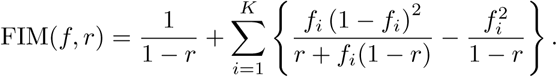

The notation FIM(*f, r*) reflects our consideration of the FIM as a function of *f* and *r*. We now wonder how to maximize FIM over the vector *f* = (*f*_1_, …, *f*_*K*_), for any *r*. This is a constrained and nonlinear optimization problem since *f* has to be made of non-negative entries and sums to one (thus *f* is in the simplex of dimension *K*). We restrict our attention to *r* ∈ (0, 1), that is *r* ≠ 0 and *r* ≠ 1, since the interpretation of FIM as a measure of the precision of the maximum likelihood estimator is only valid when *r* is away from the boundaries of the parameter space [0, 1]. For *r* ∈ (0, 1), the function *f* ↦ FIM(*f, r*) is finite and continuous, on the simplex which is a compact set, thus it attains a maximum according to the extreme value theorem.

After plotting the contours of the function FIM on the simplex and for different values of *r* (and perhaps noticing that *f* ↦ FIM(*f, r*) is symmetric with respect to the center of the simplex), we gather that the maximizer might be *f* ^*\**^ = (*K*^−1^, …, *K*^−1^), irrespective of the value of *r*. We now prove that this is indeed the case. We do so by considering an *f* such that *f*_1_ < *f*_2_. We will see that we can increase FIM(*f, r*) by modifying *f* as follows: define 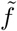 as 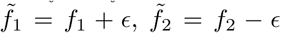 and 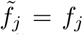 for all *j* ∈ {3, …, *K*} (if *K* ≥ 3). We will see that there exists an *ϵ* > 0 such that FIM(*f, r*) > FIM(*f, r*). Since this holds for all *f* with a pair of non-equal entries, we will be able to conclude that the unique maximizer of FIM is *f* ^*\**^ = (*K*^−1^, …, *K*^−1^).

So let us consider *f* with *f*_1_ < *f*_2_. We start by noting that, for all *f* ∈ (0, 1),

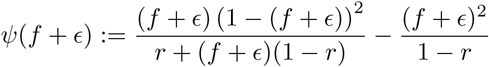

can be expanded as *ϵ* → 0 as

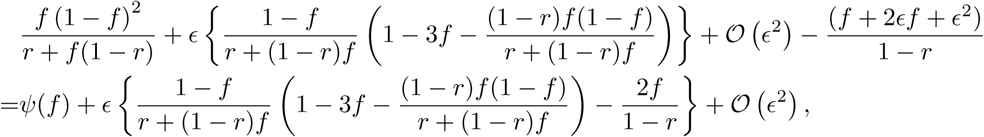

where 𝒪 (*ϵ*^2^) refers to terms which behave as *ϵ*^2^ when *ϵ* → 0 and thus are negligible in front of the term in *ϵ*. From this we deduce that *ψ*(*f* + *ϵ*) *- ψ*(*f*) = *ϵh*(*f*) + 𝒪 (*ϵ*^2^) with

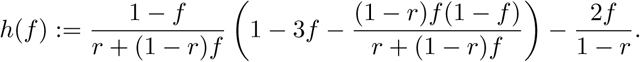

We now show that *f* ↦ *h*(*f*) is decreasing in *f* over [0, 1]. We do so by bruteforce differentiation, yielding

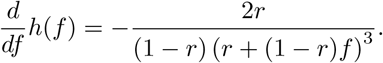

We see that the above expression is strictly negative for all *r* and *f* so that *f* ↦ *h*(*f*) is strictly decreasing.

The fact that *f* ↦ *h*(*f*) is strictly decreasing allows us to conclude the proof. Indeed, combined with the assumption *f*_1_ < *f*_2_, we have *h*(*f*_1_) > *h*(*f*_2_). Therefore,

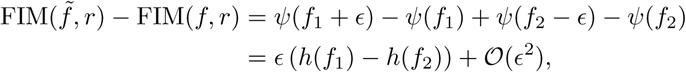

from which we deduce that there is an *ϵ* > 0 small enough so that FIM 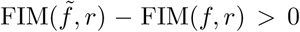. To summarize, if *f* is such that one of its components is strictly greater than another component, then we can increase the objective function FIM. We deduce that the function *f* ↦ FIM(*f, r*) is uniquely maximized at *f* ^*\**^ = (*K*^−1^, …, *K*^−1^), for which no component is greater than another one.

## Footnotes

1 IBD was first defined it terms of mutation: ‘pairs of alleles at a locus are mutation-sense IBD if there has been no mutation since their MRCA’, where MRCA stands for most recent common ancestor [2]. It can also be interpreted in terms of IBD segments: shared genomic regions unbroken by recombination since their MRCA [7, 2]. This interpretation, referred to as ‘recombination-sense’ in [2], circumvents the problem of a specifying a reference population but presents the problem of specifying some small segment length [7, 2].

2 For reasons outlined at the end of this section (3.1), we purposely omit reference to either an ancestral population or segment unbroken by recombination; IBD*t* is simply a binary variable in {0, 1}.

3 Coverage is equal to the fraction of confidence intervals that contain the data generating *r*.

